# Large-scale global molecular epidemiology of antibiotic resistance determinants in *Streptococcus pneumoniae*

**DOI:** 10.1101/2025.02.14.636407

**Authors:** Kazi Shefaul Mulk Shawrob, Achal Dhariwal, Gabriela Salvadori, Rebecca Ashley Gladstone, Roger Junges

## Abstract

**Background:** *Streptococcus pneumoniae* is a leading pathogen in terms of deaths attributable to or associated with antimicrobial resistance globally. Thus, monitoring antibiotic resistance determinants constitutes a key aspect of surveillance efforts.

**Aim:** Leveraging on publicly available whole-genome sequencing data, we aimed to investigate the presence and distribution patterns of antibiotic resistance in *S. pneumoniae* with a focus on multi-drug resistance and serotype distribution.

**Methods:** Metadata and genomes were obtained from the National Center for Biotechnology Information Pathogen Detection database. Curation and harmonization were performed in R and SPSS. Data on resistance patterns was defined according to AMRFinderPlus and a combination of prediction tools were employed for *in silico* serotyping.

**Results:** Analyses involved 75,161 genomes totaling 122,673 gene/allele counts from 14 antibiotic classes. Multi-drug resistance was observed in 16.7% of isolates with the highest increasing rates in Asia and South America. Within antibiotic classes, increase in macrolide resistance genes was highlighted, particularly in the proportion of genomes presenting *mefA*/*msrD*. Over a third of isolates with serotypes 19F, 23F, 15A, 6B, and 19A showed multi-drug resistance. We further observed the highest significant increases in the presence of resistance in 33F, 22F, 10A, and 23A. Serotype 13, not included in any vaccine formulation, presented high multi-drug resistance rates with a strong increasing trend.

**Conclusions:** The findings of this study highlight variations in resistance determinants globally and across serotypes over time. Collectively these data underscore the added value of utilizing public whole-genome sequencing data to investigate the effectiveness and repercussions of treatment and vaccination strategies on managing antibiotic resistance.

## Introduction

*Streptococcus pneumoniae* is a leading cause of noninvasive and invasive disease worldwide, especially in young children and older adults [1]. In 2019, *S. pneumoniae* was one of five pathogens – together with *Staphylococcus aureus*, *Escherichia coli*, *Klebsiella pneumoniae*, and *Pseudomonas aeruginosa* – estimated to cause over 4 million deaths worldwide. The impact was particularly severe in children as *S. pneumoniae* was the pathogen associated with most deaths among children younger than 5 years [2, 3]. The development and spread of antimicrobial resistance (AMR) create further challenges as it leads to longer hospital stays, higher mortality rates, and a substantial economic burden [4].

The capsular polysaccharide is the primary virulence determinant in *S. pneumoniae* with current vaccine strategies being based on protection against a selection of these serotypes. Understanding which serotypes have higher chances of carrying antibiotic-resistant determinants is crucial for clinical treatment as well as informed decisions on vaccine strategies employment and further research development. There are six available pneumococcal conjugate vaccines (PCVs) and one pneumococcal polysaccharide vaccine (PPSV23). These vaccines have helped to reduce the incidence of invasive disease [5]; however, coverage is a challenge [6] and less than a fourth of over 100 serotypes are currently included in these vaccines, which creates risks for the emergence of non-vaccine serotypes [7, 8]. As the pneumococcus exhibits high rates of horizontal gene transfer, further risks of serotype switching, and vaccine escape are also significant [9]. The emergence of beta-lactam resistance in *S. pneumoniae* in the 80s and 90s lead to a higher utilization of other classes of drugs such as macrolides and fluoroquinolones. Recently, the World Health Organization (WHO) has published the 2024 Bacterial Priority Pathogens List [10], which includes macrolide-resistant *S. pneumoniae*, as a response to the concerning rise in resistance in the species particularly in low- and middle-income countries (LMIC) and in vulnerable populations [11, 12]. Globally, rates of macrolide resistance vary depending on the region with a wide range of 1-99% [13–15].

Microbial phenotypical testing in the laboratory remains the gold standard for assessment of resistance patterns in pathogens to assist clinical treatment. However, the utilization of whole genome sequencing (WGS) data can enhance the surveillance efforts for AMR, as the detailed information obtained from genomic sequences allows the investigation of specific genes and their alleles, together with the integration of metadata and further virulence markers [16, 17]. This strategy allows for the understanding of genetic distribution of resistance determinants, trends in development, co-occurrence patterns, multi-drug resistance (MDR), among others [18, 19]. While phenotypic and genomic data integration challenges persist, the benefits of this combined strategy are substantial [19]. In addition, recent studies indicate strong correlations between genotypic inference and phenotypical data [20, 21]. With the further advancement of on-site sequencing speeds and machine learning tools for phenotype prediction, genomic assessment might potentially be employed to enable rapid clinical diagnosis and decision making [22, 23]. Regardless, it remains a valuable tool for retrospective analyses.

As the number of genomes available in public repositories rapidly increases, this manifests as an important resource for genomic surveillance. Recent studies have shown the added value of utilizing such data for tracking and surveillance of antimicrobial resistance in probiotic bacteria [24] and in different human and animal pathogens including *Escherichia coli* [25], *Salmonella* [26], *Campylobacter* spp. [27], *Staphylococcus aureus* [28], and *Streptococcus agalactiae* [29]. As such, the aims for this study were to leverage pneumococcal genomic data mined from public repositories to investigate distribution of ARGs, occurrence of MDR, and temporal patterns of resistance across serotypes.

## Materials and Methods

### Genomic data

Data was obtained from the National Center for Biotechnology Information (NCBI) Pathogen Browser (https://www.ncbi.nlm.nih.gov/pathogens/) on August 12^th^, 2024, which includes genomic sequences and metadata from a variety of sources including public health surveillance programs. In the database, antimicrobial resistance gene classification was performed based on the AMRFinderPlus tool (v3.11.26 and v3.12.8) [30]. Reported resistance genes or divergent alleles calling follows the reference gene catalog for the referred resource tool (https://www.ncbi.nlm.nih.gov/pathogens/refgene/). Further, antimicrobial class and subclass categories that were followed are available on Github (https://github.com/evolarjun/amr/wiki/class-subclass). For subsets of analyses, genomic sequences were retrieved utilizing the NCBI Datasets pipeline and the match data function in R was employed to obtain metadata information based on their respective accession numbers.

### Dataset curation

Data were imported into R (v4.3.2). Curation of metadata was performed with the tidyverse (v2.0.0), dplyr (v1.1.4), stringr (v1.5.1), magrittr (v2.0.3), and countrycode (v1.6.0) packages. Further details on data handling and curation are available in the supplementary material.

### Multidrug resistance and resistance determinant calling

Multidrug resistance (MDR) was defined as isolates with reported resistant genotypes to three or more unique classes. A new MDR classification column was created and a Multidrug Resistance Score (MDR Score) was calculated based on previous studies [25, 26]. MDR score measures the number of antibiotic classes a genome is predicted to be resistant to, based on antimicrobial resistance genes (ARGs) or divergent alleles. This score uses a discrete numerical scale, removing duplicates, and counting unique antimicrobial classes. For antibiotic resistance gene calling, NCBI’s AMRFinderPlus [30] criteria were employed with only complete genes being included. The only exception was that we excluded the *pmrA* gene, which composes a transporter system together with *pmrB* that is often associated with resistance to antimicrobial peptides and can also confer resistance to some antibiotics. Its presence is mostly ubiquitous in *S. pneumoniae* and its mode of function differs from conventional ARGs [31, 32], thus we opted for removal from the dataset. In addition, it is important to note that penicillin-binding proteins (*pbps*), are core genes in streptococci; however, they can exhibit divergent allele variations and combinations conferring resistance to a variety of beta-lactam antibiotics. AMRFinderPlus reports the three main *pbps* when alleles show protein identity below pre-defined thresholds of 99% for *pbp1a*, 99% for *pbp2b*, and 98% for *pbp2x*, as compared to the reference gene catalog.

### Serotype classification and statistical analyses

To investigate the relationship between ARGs and capsule in this collection of isolates, serotyping was performed by retrieving sequences for all genomes linked to an accessioned assembly (n=45,376). Details regarding the serotyping strategy are available in the supplemental material. For convenience, the list of serotypes included in each vaccine category is available in Table S1. Cross-tabulations and descriptive analyses were performed using SPSS and R Studio. A two-sided P value of <0.05 was considered statistically significant. For temporal trend analysis utilizing MDR score, negative binomial regression or Poisson regression were employed using MASS package (v7.3-60.0.1), considering count data as a metric and the overdispersion presented in the analyzed data. Temporal trend analyses with relative numbers were performed employing quasibinomial regression.

## Results

### Data description

Metadata for 75,161 *S. pneumoniae* genomes were analyzed. Of these, 53,606 genomes presented location information with continent distribution as North America (n=23,356; 31.1%), Africa (n=11,362; 15.1%), Asia (n=9,670; 12.9%), Europe (n=5,195; 6.9%), Oceania n=2,377; 3.2%), and South America (n=1646; 2.2%). Figure 1A shows a global heatmap of number of isolates per country, supplemented by Table S2. Year of collection was available for 51,864 genomes and their distribution is available in Figure 1B. Source of collection was most commonly reported as sterile site (n= 17,642; 23,5%) and blood (n= 10,945; 14,6 %). Missing information was commonly found in the metadata analyses, as 21,555 (28.7%) presented no location information, 23,297 (31.0%) presented no date for collection, and 26,392 (35.1%) listed no isolation source.

**Fig 1.**
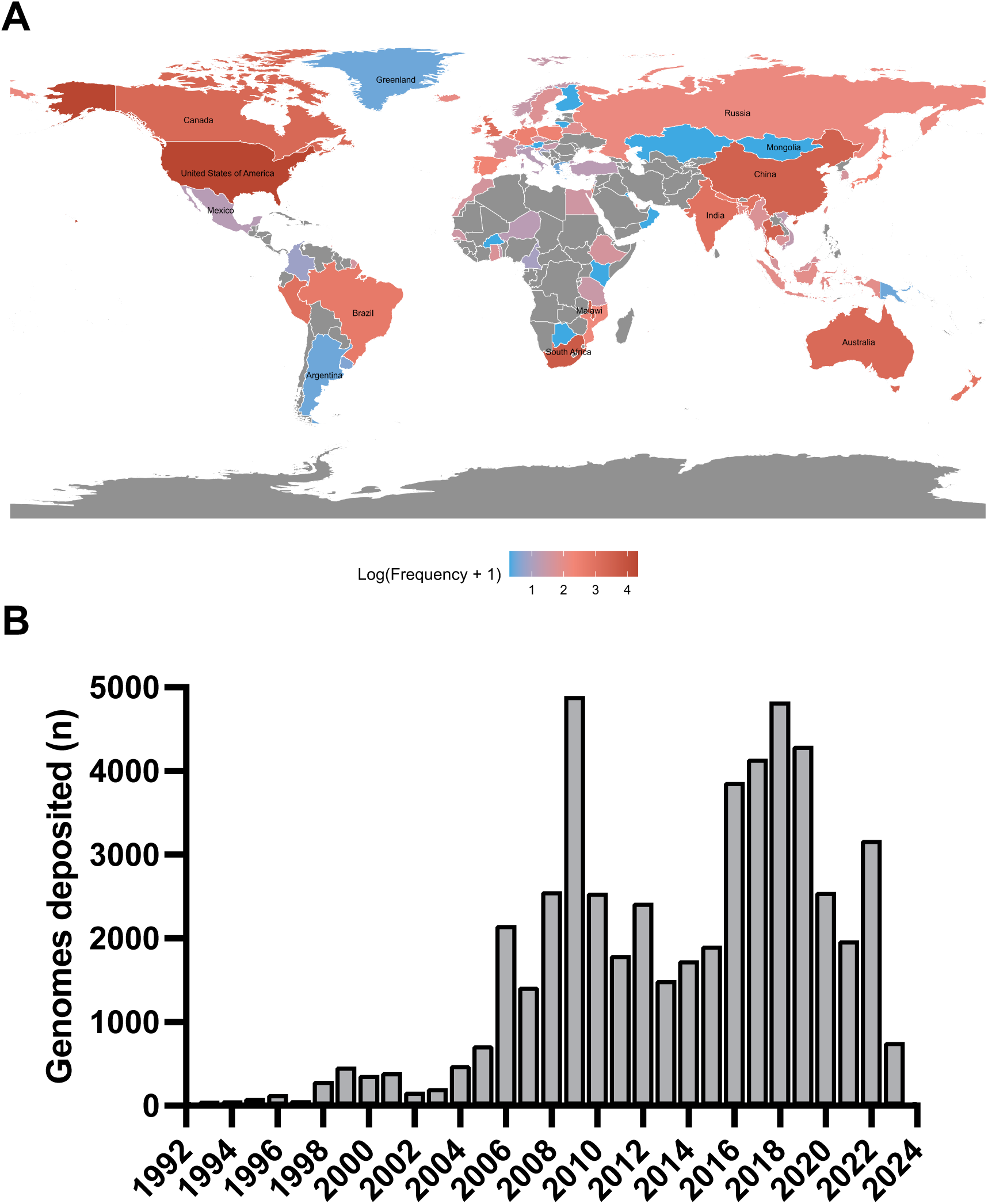
Geographic distribution and year of collection for *S. pneumoniae* isolates. (A) Global heatmap of number of isolates per country in logarithmic scale. (B) Year of collection with data shown from 1992 and forward as >40 genomes have been consistently collected since then. Prior to 1992, a total of 209 genomes distributed across ten decades were collected.

### Multi-drug resistance and gene co-occurrence

In total, 12,554 genomes (16.7%) presented MDR predicted by the genotype. As such, the average MDR score across all samples was 0.98 with a standard error of the mean of 0.004. Most genomes presented no ARGs (n=35,695) and the maximum resistance determinants observed were in three genomes with resistance to seven antibiotic classes. Distribution over years remained stable considering all world regions. Figure 2A shows MDR proportion over the years, while Figure 2B reveals a declining trend in MDR scores over time. Data for each country and region on MDR is available in Figure 2C. With data stratified by world region, the strongest increasing trends were observed for Asia and South America in recent years in terms of MDR score (Figure 3). In MDR isolates, the association between divergent-*pbp2bs* and *tet(M)* showed the highest frequency (10,943 co-occurrences). Additionally, *erm(B)* and *tet(M)* co-occurred 9,109 times, while *erm(B)* and divergent-*pbp2bs* were found together in 8,346 instances (Figure 2D).

**Fig 2.**
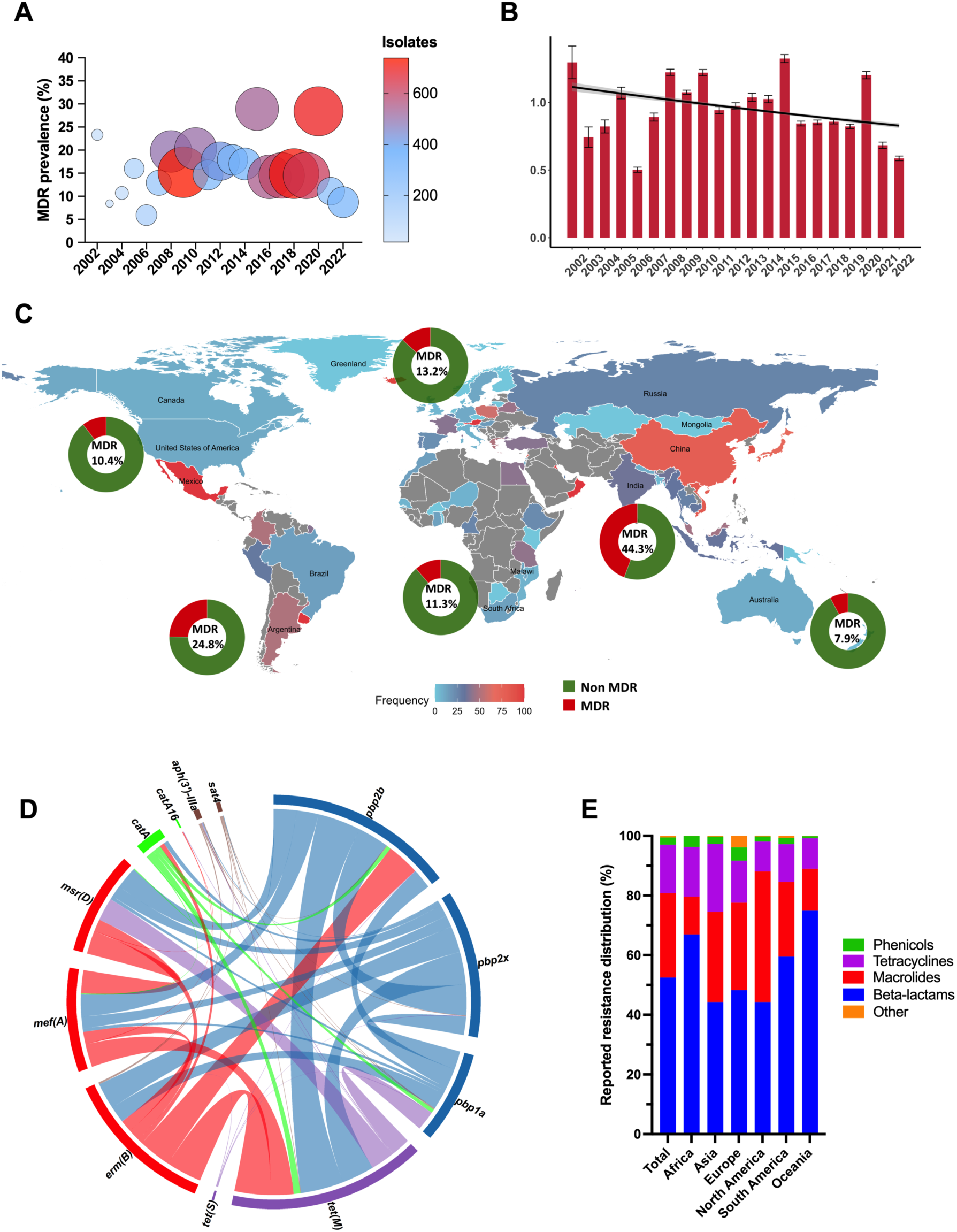
Presence of multidrug resistance in *S. pneumoniae* isolates. (A) Proportion and number of isolates with MDR by year. (B) MDR score over time using negative binomial regression (p<0.05). (C) Relative presence of MDR per continent in each donut plot and by country according to the color code. (D) Chord diagram of gene co-occurrence in MDR isolates (threshold of 100 occurrences and above), with genes from the same antibiotic class grouped into one color. (E) Distribution of resistance towards the four most frequent classes in the total population and in each continent.

**Fig 3.**
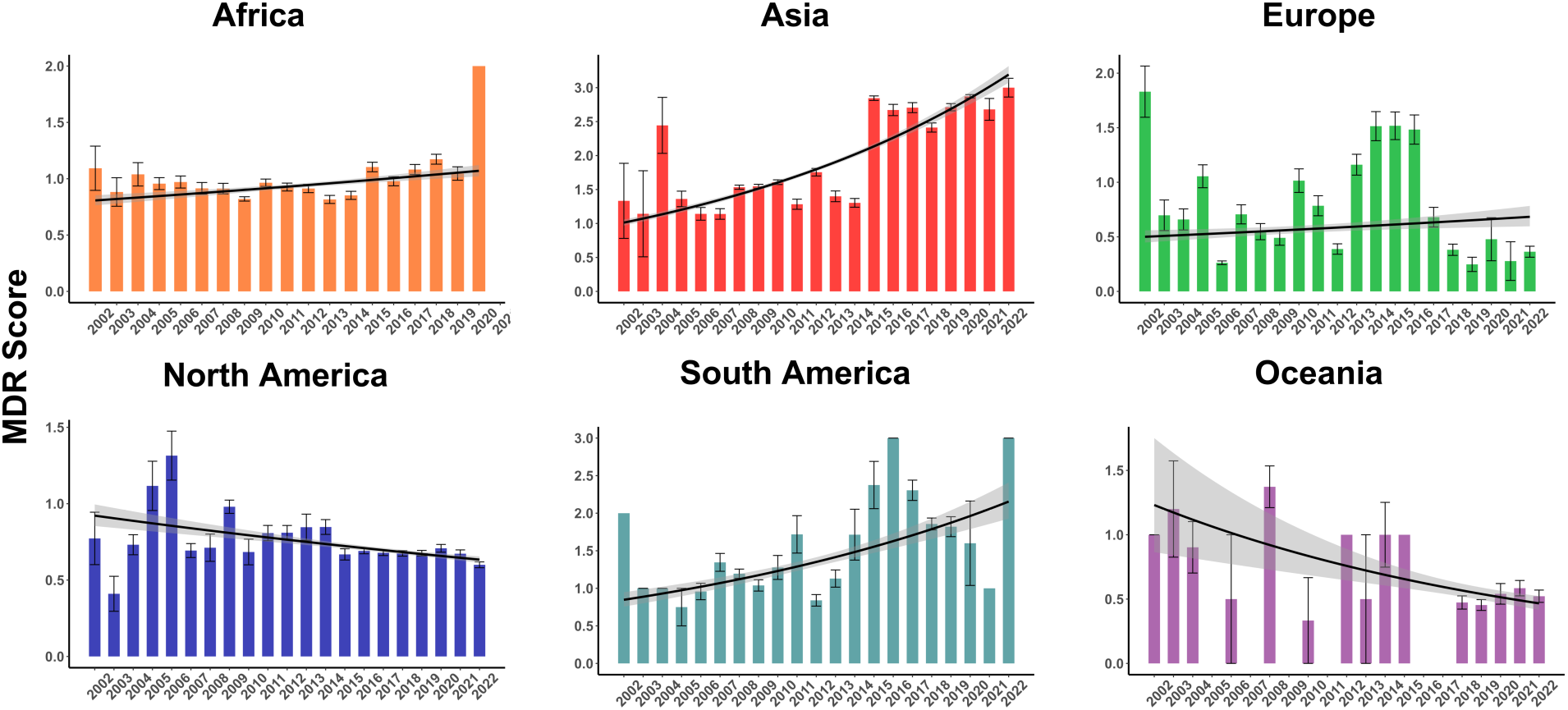
MDR score by year in each continent. Bars represent mean MDR scores with SEM, and lines represent the fitted MDR scores derived from negative binomial regression, while the shading the 95% confidence intervals. All trends were significant (***, p < 0.001), with the exception of Europe, where a lower significance level was observed (*, p < 0.05).

### Class and gene distribution

A total 72 unique resistance determinants and 122,673 counts of either ARGs or reported resistance alleles were observed. The four antibiotic classes in which resistance was commonly predicted to were beta-lactams (64,427; 52.5% of the total ARGs and 40.9% of genomes containing at least one ARG/divergent allele), macrolides (34,641; 28.2% of the total ARGs and 25.9% of genomes containing at least one ARG), tetracyclines (19,884; 16.2% of the total ARGs and 26.4% of genomes containing at least one ARG), and phenicols (2,986; 2.4% of the total ARGs and 4.0% of genomes containing at least one ARG). Relative distribution of these classes in total and in each continent is available in Figure 2E. Development trends over the years for the three most common classes in each continent are shown in Figure S1. The list of ARGs/divergent resistance alleles identified is available in Table S3 and their distribution across continents as a heatmap is shown in Figure 4. For the eight most frequent resistance determinants, temporal development is visualized in Figure 5. Over time, presence of determinants that can confer resistance to beta-lactams and tetracyclines is seemingly decreasing while an increase in macrolide resistance was observed. For genotype distribution regarding macrolide resistance, higher proportions of *mef(A)* resistance were observed (16%, 42% of genomes) compared to *erm(B)* (14.3%), with 4.2% of genomes presenting dual-genotype resistance. However, this varied across world regions (Table S4). While genomes from Africa and North America showed predominance of *mef(A)* genotype, the opposite was seen for isolates from Asia, Europe, South America, and Oceania. For less frequently present genes, *tet32* stood out with 160 occurrences given that a significant number has been observed from 2017-2022 (n=67).

**Fig 4.**
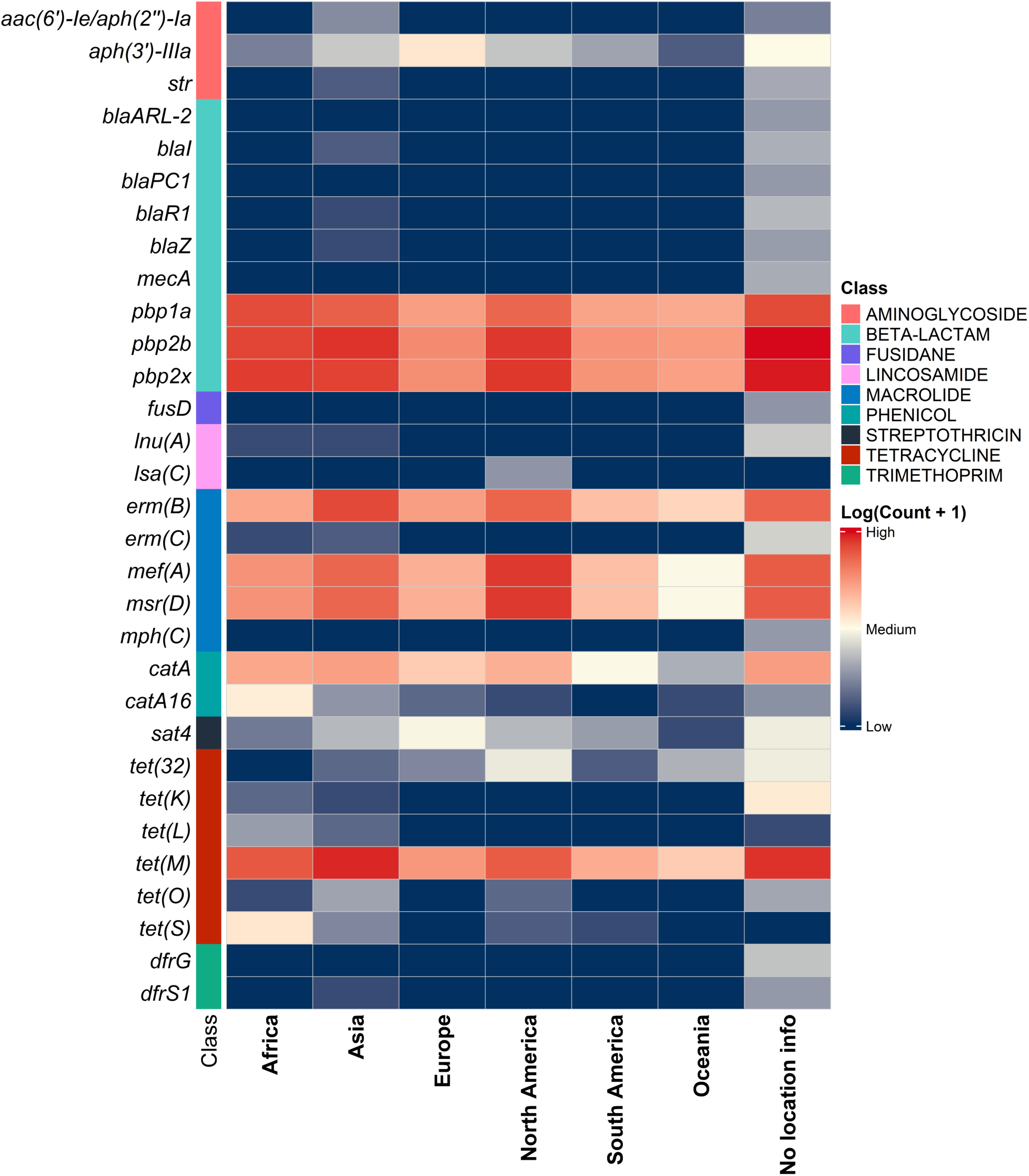
Heatmap showing relative presence of resistance genes and divergent *pbp*s in *S. pneumoniae* genomes grouped by continent. Genes are grouped by color according to antibiotic class and genes with a count below 10 were removed. The complete list of genes is available in Table S4.

**Fig 5.**
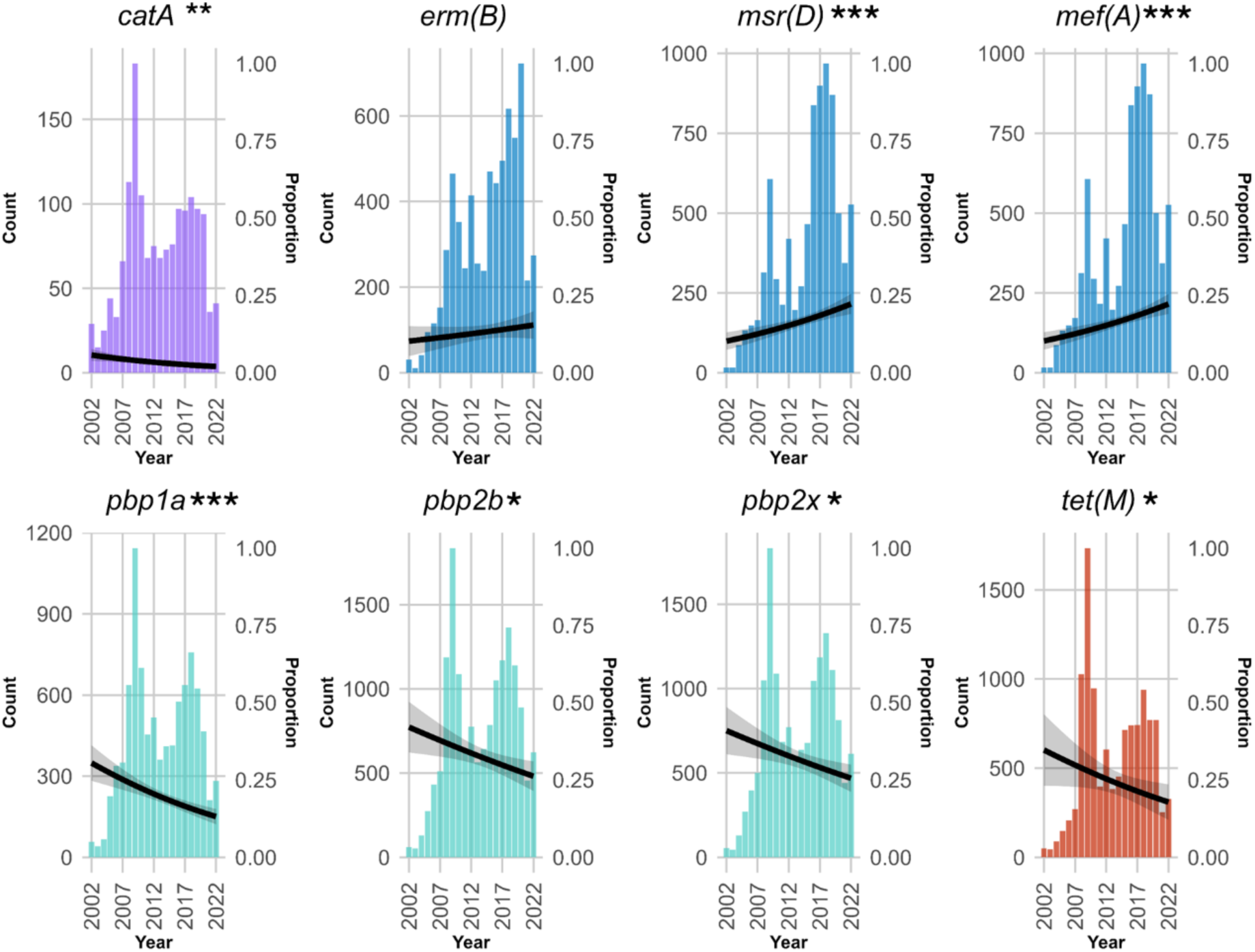
Development over time for the eight most common resistance determinants in *S. pneumoniae* isolates. Colors indicate grouping by antibiotic class. Lines represent the fitted MDR proportion derived from quasibinomial regression, while the shading the 95% confidence intervals. Fitted curves were generated with quasi-binomial regression analysis, also indicating statistical significance. *p<.05, **p<0.01***p<.001

### Serotype association with AMR

In total, 40,005 sequences had their serotypes predicted. Analysis of most common serotype and their distribution over time is shown in Figure S2. The 5 serotypes with higher proportion of MDR were 19F (63.2%), 15A (43.3%), 23F (43.3%), 6B (42.5%%), and 19A (34.8%). List of the 25 most prevalent serotypes with genome counts and MDR rates are available in Table S5. When looking at distribution of the MDR score over time, several serotypes presented different trends of resistance determinants accumulation (Figure 6A). The higher significant increases were seen in 33F, followed by 22F, 10A, 23A, and 35B. In comparison, serotypes 34, 19F, and 16F showed declining patterns. Individual graphs for each serotype are shown in Figure S3. Figure S4 shows relative predicted resistance to each serotype to the most common antibiotic classes. Among serotypes not included in any vaccine formulation, serotype 13 presented the highest MDR percentage at 11.7%, followed by 6C at 7.5% and 7C at 6.3% (Table S6). When comparing serotypes included or not in PCV13, we observed a sharper increase in the accumulation of resistance determinants in non-vaccine serotypes (Fig 6B). This was also observed for all the other vaccine formulations available (data not shown).

**Fig 6.**
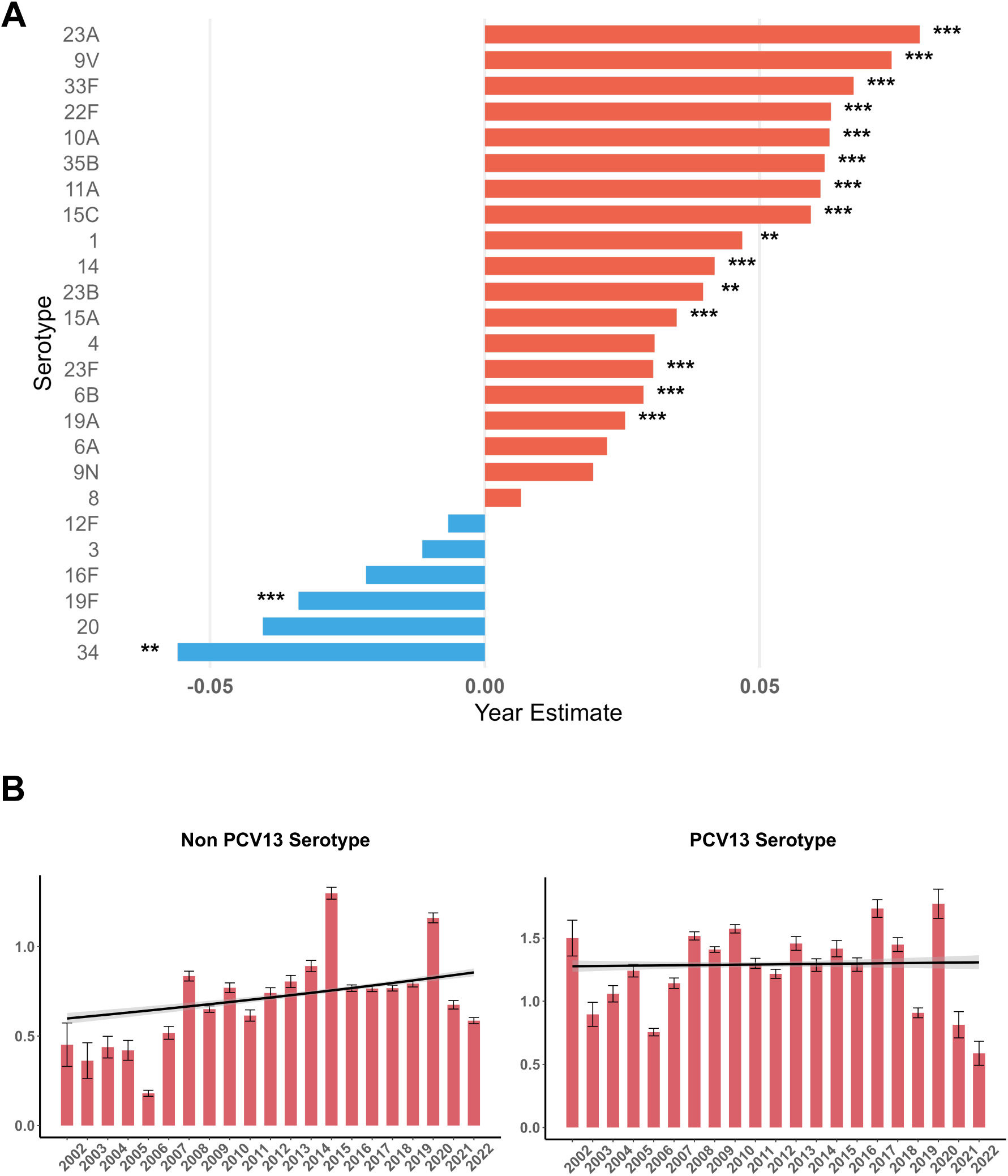
Development trends of MDR score. (A) Across serotypes of *S. pneumoniae* from 2002 to 2022 with over 500 genomes available in the dataset. Bars represent estimates of yearly changes in MDR score per serotype. Asterisks indicate statistical significance by Poisson or negative binomial regression (B) Across serotypes included and not included in PCV13. Bars represent the mean MDR scores with SEM, while lines indicate the fitted MDR scores from negative binomial regression, with shading representing the 95% confidence intervals. For non PCV13 serotypes, the estimated yearly change in MDR score is 0.0178 (***), while for PCV13 serotypes, the estimated yearly change in MDR score is 0.0012 (ns).

## Discussion

Analyses including data from over 75,000 genomes showed an average of just under one reported resistance determinant per pneumococcal genome coupled with MDR being identified for 16.7% of genomes. Stable numbers of MDR were observed globally over the years; however, data from genomes collected from Asia and South America indicate strongly increasing trends in MDR. Of particular concern was the increase of macrolide resistance determinants. We utilized an *in-silico* approach to predict the serotype for about 40,000 genomes and identified a high number of resistance determinants in serotypes 19F, 23F, 15A, 6B, and 19A. Further, upward trends of resistance presence were observed for a number of serotypes such as 33F, 22F, 10A, and 23A; while decreasing trends were seen for 34, 19F, and 16F. Non-vaccine serotype 13 presented a considerable MDR rate and a sharp increasing trend in ARG presence, which warrants monitoring.

Multidrug-resistant *S. pneumoniae* in young children is a significant concern [33]. We identified an overall average rate of MDR around 16%. Specific rates for each region and country varied significantly. Different study and sampling strategies make it likely that some over- and under-estimations are present in the analyzed dataset. However, the trend of high prevalence of MDR in Asia aligns with previous research that has reported elevated MDR rates in *S. pneumoniae* isolates, with recent data indicating rates of up to 92% [34–36]. High rates of MDR and macrolide resistance in South America have also been described previously [11, 37]. The South America region has a low number of sequences collected (n=1,562). This is coupled with an MDR rate of over 24% and an increasing trend in recent years and highlights the necessity for increased surveillance efforts as an increase in antibiotic usage in both South America and Asia is likely to be seen in upcoming years, together with rapid economic development in parts of these continents [38].

Macrolide resistance is a growing concern for pneumococci mediated by two main mechanisms. As macrolides target protein synthesis, *erm*(B) acts as a ribosomal methylase that dimethylates the 23S rRNA, preventing binding of the drug. Often the phenotype observed is resistance to macrolide-lincosamide-streptograminB, referred to as MLSB phenotype [39]. On the other hand, *mef*(A/E) works as an efflux pump in tandem with *msr(D)* by removing the drug from the cellular environment. This M phenotype confers resistance to 14- and 15- membered macrolides but is sensitive to clindamycin, in contrast with the MLSB phenotype [40]. Distribution of each ARG varies and the widespread presence of genomes containing both genes has been highlighted as a concern [12, 41–44]. Our study identified *erm(B)* as the most dominant gene responsible for macrolide resistance in Asia, Europe, and Oceania whereas *mefA*/*msrD* was most prevalent in North America and in Africa. These patterns have been observed previously in a variety of studies (for review see [12, 41]). We also show that the rise in macrolide resistance is seemingly driven by *mefA/msrD*. A significant proportion in dual genotype (*ermB* and *mefA*/*msrD*) macrolide resistance was noted in Asia and South America with rates over 20%, while in other regions rates it ranged between 1-5%. Isolates with dual genotype have been shown to present clinical resistance to macrolides similar to isolates presenting only *erm(B)* [43]. Therefore, while presence of two redundant ARGs does not seem to lead to higher level of macrolide resistance, these isolates have been often associated with more MDR phenotypes. We observed in our study that 98% of dual genotype isolates were also MDR and 97.3% presented resistance to beta-lactams, in comparison to 88.7% / 84.9% in pure *erm(B)* isolates and 49.1% / 69.7% in *mef(A)* and *msr(D)* isolates. A plausible explanation for the finding can be the presence of multiple mobile genetic elements in dual genotypes isolates. The *erm(B)* gene is often associated with integrative and conjugative elements (ICEs) such as Tn*6002*, Tn*6003*, Tn*6009* and Tn*1545*, that often co-carry other ARGs such as *tet(M)* and *aphA-3* [45–47]. Additionally, the mega element in *S. pneumoniae* carries the *mef(E)* variant of *mef(A)*, and it is often also present with other resistance genes in Tn*2009*, Tn*2010*, and Tn*2017* [48–50]. Together, the presence of two possible ICEs coupled with the high proportion of penicillin-binding-proteins in the species increases the chance of MDR frequency in these isolates.

The two most common serotypes observed in this study were 19F and 19A, and both were associated with high rates of MDR. These serotypes are often involved with invasive pneumococcal disease (IPD) in children and adults [51–59]. However, their involvement with IPD has decreased post-PCV introduction [58, 60]. From 2002 to 2022, we observed a decrease in resistance proportion in 19F isolates, which coincides with its decrease in involvement with IPD globally. On the other hand, serotype 19A emerged after introduction of PCV7 [61]. Following the inclusion in PCV13, its rates of involvement with IPD disease have gone down [62, 63]. In Belgium, after switching from PCV-13 to PCV-10, a rapid reemergence in serotype 19A causing IPD particularly in children was observed [64]. Further, it has been shown that serotypes 19A, 3, and 7C persist as nasopharyngeal colonizers even post introduction of PCV13 [65]. Looking at development over time, an upwards trend was observed for serotype 19A between 2002-2022, mainly driven by the period pre-PCV13. For serotype 3, another colonizer with high virulence trains (for review see [66]), our results indicate stable levels of determinants over the years, but an increase in proportion of genomes containing *tet32*, a tetracycline resistance gene, calls for attention and should be further monitored moving forward. The successive accumulation of ARGs even in serotypes that are covered by vaccine strategies is a concern given how prolific streptococci are at exchanging genes via horizontal gene transfer.

Upwards trends of ARG/divergent allele presence were observed for a variety of non-vaccine serotypes such as 23A and 23B. While these strains are generally more often associated with carriage and less associated with IPD [67, 68], the emergence of resistance is cause of concern particularly for populations at higher risk of pneumococcal disease. Cases of disease for 23A and 23B have also been observed more frequently after introduction of PCV13 [63], as well as higher levels of resistance [69, 70]. 23A and 23B have only been included in the recent PCV21 which tries to tackle problematic serotypes in adults. Serotype 13 is not included in vaccine formulations currently and has shown a significant rise in resistance determinants recently. The introduction of PCVs has clearly reduced pneumococcal disease rates globally [5] and has had positive general effects on AMR [71]. However, due to limitations in the number of serotypes that are covered in the PCVs, their implementation has led to significant changes in serotype distribution, with non-vaccine types increasing in proportion [72, 73]. Understanding these influences is central for interpreting the relationship between vaccine implementation and serotype diversity and will continue to be crucial to follow up on population dynamics moving forward.

While this study provides a global overview of AMR in *Streptococcus pneumoniae* isolates, it is important to acknowledge several limitations in the dataset. Data available in public repositories are not the result of systematic random sampling and can present reporting bias resulting in over- or under-presenting AMR, thus we have presented trends and highlight potential threats to the future control of AMR in pneumococci. Generally, recent studies have shown a high level of accuracy regarding *in silico* prediction of AMR [30, 74, 75] and serotype [76–78]. In addition, it is likely that a proportion of the predicted resistance determinants will be associated with no or intermediate resistance, *e.g.* divergent *pbps* might not necessarily lead to consistent and predictable beta-lactam resistance. In such instance, mosaic sequences and combination of *pbp* mutations are challenging to predict and classify [79], including also the involvement of additional genetic elements such as the *murMN* operon that encodes enzymes involved in cell wall synthesis [80]. The combination of such factors may lead to various levels of penicillin resistance and extended-spectrum cephalosporin resistance [81, 82]. In our analyses, a decrease in divergent *pbp* proportion was observed, so it is unlikely that possible overestimations of predicted *pbp* phenotypes have impacted trend analyses for MDR and serotypes. It is important to highlight, however, that further research on this topic including sequences from closely related commensal species is warranted to better unravel the puzzle of beta-lactam resistance in streptococci from the mitis group. Nevertheless, timely detection of increased spread of resistance determinants can offer an early warning system to surveillance authorities and guide future research.

The sharing and reusability potential of genomic data is a priority highlighted by a number of stakeholders worldwide [18, 83, 84]. An important advantage of WGS monitoring is the possibility of linkage with metadata. This allows for the comparison across collection sites coupled with the possibility of correlation with clinical characteristics. Thus, poorly curated or lack of metadata significantly impair the potential for such analyses. In the data utilized in this study, over a third of genomes lacked information on time of collection, location, and source. These aspects are likely to play a major role moving forward as NGS technologies become more affordable and the collective amount of WGS-data rapidly increases. However, to fully explore the potential of public repositories for genomic data, it is crucial that better standardization pipelines for depositing genomic data are adopted [85, 86].

This study provides valuable insights for tracking pneumococcal resistance trends and highlights the importance of continuous surveillance. Such findings can assist in guiding the development of effective policies aimed at managing pneumococcal resistance and ensuring appropriate interventions. We highlight variations in the presence of resistance determinants globally as well as across serotypes over time in *S. pneumoniae*. Collectively, these data underscore the added value of utilizing public data to gain knowledge on the effectiveness and repercussions of treatment and vaccination strategies.

## Data availability statement

Data utilized in this study are public and available in the NCBI Pathogen Detection Isolates Browser (https://www.ncbi.nlm.nih.gov/pathogens/) accessed on Aug 12^th^, 2024.

## Author contributions

Study concept and design: KS and RJ; acquisition of data: KS; analysis and interpretation of data: KS, AD, GS, RG, and RJ; drafting of the manuscript: KS and RJ; critical revision of the manuscript: KS, AD, GS, RG, and RJ; statistical analysis: KS; study supervision: RJ. All authors read and approved the final manuscript.

## References

1. Weiser JN, Ferreira DM, Paton JC. Streptococcus pneumoniae: transmission, colonization and invasion. Nat Rev Microbiol. 2018;16(6):355–67.

2. O’Brien KL, Wolfson LJ, Watt JP, Henkle E, Deloria-Knoll M, McCall N, et al. Burden of disease caused by Streptococcus pneumoniae in children younger than 5 years: global estimates. Lancet. 2009;374(9693):893–902.

3. G. B. D. Antimicrobial Resistance Collaborators. Global mortality associated with 33 bacterial pathogens in 2019: a systematic analysis for the Global Burden of Disease Study 2019. Lancet. 2022;400(10369):2221–48.

4. Antimicrobial Resistance C. Global burden of bacterial antimicrobial resistance in 2019: a systematic analysis. Lancet. 2022;399(10325):629–55.

5. Wahl B, O’Brien KL, Greenbaum A, Majumder A, Liu L, Chu Y, et al. Burden of Streptococcus pneumoniae and Haemophilus influenzae type b disease in children in the era of conjugate vaccines: global, regional, and national estimates for 2000-15. Lancet Glob Health. 2018;6(7):e744–e57.

6. Kim C, Holm M, Frost I, Hasso-Agopsowicz M, Abbas K. Global and regional burden of attributable and associated bacterial antimicrobial resistance avertable by vaccination: modelling study. BMJ Glob Health. 2023;8(7).

7. Dagan R. Impact of pneumococcal conjugate vaccine on infections caused by antibiotic-resistant Streptococcus pneumoniae. Clin Microbiol Infect. 2009;15 Suppl 3:16–20.

8. Lo SW, Gladstone RA, van Tonder AJ, Lees JA, du Plessis M, Benisty R, et al. Pneumococcal lineages associated with serotype replacement and antibiotic resistance in childhood invasive pneumococcal disease in the post-PCV13 era: an international whole-genome sequencing study. Lancet Infect Dis. 2019;19(7):759–69.

9. Wyres KL, Lambertsen LM, Croucher NJ, McGee L, von Gottberg A, Linares J, et al. Pneumococcal capsular switching: a historical perspective. J Infect Dis. 2013;207(3):439–49.

10. WHO. WHO Bacterial Priority Pathogens List 2024 Geneva: World Health Organization; 2024. Available from: https://public.ebookcentral.proquest.com/choice/PublicFullRecord.aspx?p=31361074.

11. Gonzales BE, Mercado EH, Pinedo-Bardales M, Hinostroza N, Campos F, Chaparro E, et al. Increase of Macrolide-Resistance in Streptococcus pneumoniae Strains After the Introduction of the 13-Valent Pneumococcal Conjugate Vaccine in Lima, Peru. Front Cell Infect Microbiol. 2022;12:866186.

12. Schroeder MR, Stephens DS. Macrolide Resistance in Streptococcus pneumoniae. Front Cell Infect Microbiol. 2016;6:98.

13. Zhou X, Liu J, Zhang Z, Cui B, Wang Y, Zhang Y, et al. Characterization of Streptococcus pneumoniae Macrolide Resistance and Its Mechanism in Northeast China over a 20-Year Period. Microbiol Spectr. 2022;10(5):e0054622.

14. Farrell DJ, Couturier C, Hryniewicz W. Distribution and antibacterial susceptibility of macrolide resistance genotypes in Streptococcus pneumoniae: PROTEKT Year 5 (2003-2004). Int J Antimicrob Agents. 2008;31(3):245–9.

15. Debess Magnussen M, Erlendsdottir H, Gaini S, Gudnason T, Kristinsson KG. Streptococcus pneumoniae: Antimicrobial Resistance and Serotypes of Strains Carried by Children and Causing Invasive Disease in the Faroe Islands. Microb Drug Resist. 2018;24(10):1507–12.

16. Baker S, Thomson N, Weill F-X, Holt KE. Genomic insights into the emergence and spread of antimicrobial-resistant bacterial pathogens. Science. 2018;360(6390):733–8.

17. Djordjevic SP, Jarocki VM, Seemann T, Cummins ML, Watt AE, Drigo B, et al. Genomic surveillance for antimicrobial resistance—a One Health perspective. Nature Reviews Genetics. 2024;25(2):142–57.

18. Hendriksen RS, Bortolaia V, Tate H, Tyson GH, Aarestrup FM, McDermott PF. Using Genomics to Track Global Antimicrobial Resistance. Front Public Health. 2019;7:242.

19. Jauneikaite E, Baker KS, Nunn JG, Midega JT, Hsu LY, Singh SR, et al. Genomics for antimicrobial resistance surveillance to support infection prevention and control in health-care facilities. The Lancet Microbe. 2023;4(12):e1040–e6.

20. Bharat A, Petkau A, Avery BP, Chen JC, Folster JP, Carson CA, et al. Correlation between phenotypic and in silico detection of antimicrobial resistance in Salmonella enterica in Canada using Staramr. Microorganisms. 2022;10(2):292.

21. Sia CM, Baines SL, Valcanis M, Lee DY, Gonçalves da Silva A, Ballard SA, et al. Genomic diversity of antimicrobial resistance in non-typhoidal Salmonella in Victoria, Australia. Microbial Genomics. 2021;7(12):000725.

22. Eyre DW, De Silva D, Cole K, Peters J, Cole MJ, Grad YH, et al. WGS to predict antibiotic MICs for Neisseria gonorrhoeae. J Antimicrob Chemother. 2017;72(7):1937–47.

23. Nguyen M, Long SW, McDermott PF, Olsen RJ, Olson R, Stevens RL, et al. Using Machine Learning To Predict Antimicrobial MICs and Associated Genomic Features for Nontyphoidal Salmonella. J Clin Microbiol. 2019;57(2).

24. Tóth AG, Judge MF, Nagy SA, Papp M, Solymosi N. A survey on antimicrobial resistance genes of frequently used probiotic bacteria, 1901 to 2022. Eurosurveillance. 2023;28(14).

25. Pires J, Huisman JS, Bonhoeffer S, Van Boeckel TP. Increase in antimicrobial resistance in food animals between 1980 and 2018 assessed using genomes from public databases. J Antimicrob Chemoth. 2022;77(3):646–55.

26. Pires J, Huisman JS, Bonhoeffer S, Van Boeckel TP. Multidrug Resistance Dynamics in Food Animals in the United States: An Analysis of Genomes from Public Databases. Microbiol Spectr. 2021;9(2).

27. Cobo-Díaz JF, del Río PG, Alvarez-Ordóñez A. Whole Resistome Analysis in *Campylobacter jejuni* and *C. coli* Genomes Available in Public Repositories. Frontiers in Microbiology. 2021;12.

28. Zaghen F, Sora VM, Meroni G, Laterza G, Martino PA, Soggiu A, et al. Epidemiology of Antimicrobial Resistance Genes in Staphyloccocus aureus Isolates from a Public Database in a One Health Perspective-Sample Characteristics and Isolates’ Sources. Antibiotics (Basel). 2023;12(7).

29. Meroni G, Sora VM, Martino PA, Sbernini A, Laterza G, Zaghen F, et al. Epidemiology of Antimicrobial Resistance Genes in Sequences from a Public Database in a One Health Perspective. Antibiotics-Basel. 2022;11(9).

30. Feldgarden M, Brover V, Haft DH, Prasad AB, Slotta DJ, Tolstoy I, et al. Validating the AMRFinder Tool and Resistance Gene Database by Using Antimicrobial Resistance Genotype-Phenotype Correlations in a Collection of Isolates. Antimicrob Agents Chemother. 2019;63(11).

31. Gill MJ, Brenwald NP, Wise R. Identification of an Efflux Pump Gene, *pmrA*, associated with Fluoroquinolone Resistance in *Streptococcus pneumoniae*. Antimicrob Agents Ch. 1999;43(1):187–9.

32. Chen HD, Groisman EA. The Biology of the PmrA/PmrB Two-Component System: The Major Regulator of Lipopolysaccharide Modifications. Annu Rev Microbiol. 2013;67:83–112.

33. Patil S, Chen H, Lopes BS, Liu S, Wen F. Multidrug-resistant Streptococcus pneumoniae in young children. The Lancet Microbe. 2023;4(2):e69.

34. Kim SH, Song J-H, Chung DR, Thamlikitkul V, Yang Y, Wang H, et al. Changing trends in antimicrobial resistance and serotypes of Streptococcus pneumoniae isolates in Asian countries: an Asian Network for Surveillance of Resistant Pathogens (ANSORP) study. Antimicrob Agents Ch. 2012;56(3):1418–26.

35. Miao C, Yan Z, Chen C, Kuang L, Ao K, Li Y, et al. Serotype, antibiotic susceptibility and whole-genome characterization of Streptococcus pneumoniae in all age groups living in Southwest China during 2018– 2022. Frontiers in Microbiology. 2024;15:1342839.

36. Shi X, Patil S, Wang Q, Liu Z, Zhu C, Wang H, et al. Prevalence and resistance characteristics of multidrug-resistant Streptococcus pneumoniae isolated from the respiratory tracts of hospitalized children in Shenzhen, China. Frontiers in Cellular and Infection Microbiology. 2024;13:1332472.

37. Valenzuela MT, de Quadros C. Antibiotic resistance in Latin America: a cause for alarm. Vaccine. 2009;27 Suppl 3:C25–8.

38. Sader HS, Mendes RE, Le J, Denys G, Flamm RK, Jones RN, editors. Antimicrobial susceptibility of Streptococcus pneumoniae from North America, Europe, Latin America, and the Asia-Pacific region: results from 20 years of the SENTRY antimicrobial surveillance program (1997–2016). Open forum infectious diseases; 2019: Oxford University Press US.

39. Leclercq R. Mechanisms of resistance to macrolides and lincosamides: Nature of the resistance elements and their clinical implications. Clin Infect Dis. 2002;34(4):482–92.

40. Sutcliffe J, Tait-Kamradt A, Wondrack L. Streptococcus pneumoniae and Streptococcus pyogenes resistant to macrolides but sensitive to clindamycin: a common resistance pattern mediated by an efflux system. Antimicrob Agents Chemother. 1996;40(8):1817–24.

41. Linares J, Ardanuy C, Pallares R, Fenoll A. Changes in antimicrobial resistance, serotypes and genotypes in Streptococcus pneumoniae over a 30-year period. Clin Microbiol Infect. 2010;16(5):402–10.

42. Reinert RR, Filimonova OY, Al-Lahham A, Grudinina SA, Ilina EN, Weigel LM, et al. Mechanisms of macrolide resistance among isolates from Russia. Antimicrob Agents Ch. 2008;52(6):2260–2.

43. Farrell DJ, Jenkins SG, Brown SD, Patel M, Lavin BS, Klugman KP. Emergence and spread of Streptococcus pneumoniae with erm(B) and mef(A) resistance. Emerg Infect Dis. 2005;11(6):851–8.

44. Harimaya A, Yokota S, Sato K, Yamazaki N, Himi T, Fujii N. High prevalence of erythromycin resistance and macrolide-resistance genes, mefA and ermB, in Streptococcus pneumoniae isolates from the upper respiratory tracts of children in the Sapporo district, Japan. J Infect Chemother. 2007;13(4):219–23.

45. Courvalin P, Carlier C. Tn1545: a conjugative shuttle transposon. Mol Gen Genet. 1987;206(2):259–64.

46. Soge OO, Beck NK, White TM, No DB, Roberts MC. A novel transposon, Tn6009, composed of a Tn916 element linked with a Staphylococcus aureus mer operon. J Antimicrob Chemother. 2008;62(4):674–80.

47. Rice LB. Tn916 family conjugative transposons and dissemination of antimicrobial resistance determinants. Antimicrob Agents Chemother. 1998;42(8):1871–7.

48. Zhou W, Yao K, Zhang G, Yang Y, Li Y, Lv Y, et al. Mechanism for transfer of transposon Tn2010 carrying macrolide resistance genes in Streptococcus pneumoniae and its effects on genome evolution. J Antimicrob Chemother. 2014;69(6):1470–3.

49. Del Grosso M, Camilli R, Iannelli F, Pozzi G, Pantosti A. The mef(E)-carrying genetic element (mega) of Streptococcus pneumoniae: insertion sites and association with other genetic elements. Antimicrob Agents Chemother. 2006;50(10):3361–6.

50. Del Grosso M, Scotto d’Abusco A, Iannelli F, Pozzi G, Pantosti A. Tn2009, a Tn916-like element containing mef(E) in Streptococcus pneumoniae. Antimicrob Agents Chemother. 2004;48(6):2037–42.

51. Balsells E, Guillot L, Nair H, Kyaw MH. Serotype distribution of Streptococcus pneumoniae causing invasive disease in children in the post-PCV era: A systematic review and meta-analysis. Plos One. 2017;12(5):e0177113.

52. Jaiswal N, Singh M, Das RR, Jindal I, Agarwal A, Thumburu KK, et al. Distribution of serotypes, vaccine coverage, and antimicrobial susceptibility pattern of Streptococcus pneumoniae in children living in SAARC countries: a systematic review. Plos One. 2014;9(9):e108617.

53. Jayaraman Y, Veeraraghavan B, Purushothaman GKC, Sukumar B, Kangusamy B, Kapoor AN, et al. Burden of bacterial meningitis in India: Preliminary data from a hospital based sentinel surveillance network. Plos One. 2018;13(5).

54. Phongsamart W, Srifeungfung S, Chatsuwan T, Rungnobhakhun P, Maleesatharn A, Chokephaibulkit K. Causing Invasive Diseases in Children and Adults in Central Thailand, 2012-2016. Vaccines-Basel. 2022;10(8).

55. Toizumi M, Satoh C, Quilty BJ, Nguyen HAT, Madaniyazi L, Le LT, et al. Effect of pneumococcal conjugate vaccine on prevalence of otitis media with effusion among children in Vietnam. Vaccine. 2022;40(36):5366–75.

56. Hackel M, Lascols C, Bouchillon S, Hilton B, Morgenstern D, Purdy J. Serotype prevalence and antibiotic resistance in clinical isolates among global populations. Vaccine. 2013;31(42):4881–7.

57. Bertran M, D’Aeth JC, Abdullahi F, Eletu S, Andrews NJ, Ramsay ME, et al. Invasive pneumococcal disease 3 years after introduction of a reduced 1+ 1 infant 13-valent pneumococcal conjugate vaccine immunisation schedule in England: a prospective national observational surveillance study. The Lancet Infectious Diseases. 2024;24(5):546–56.

58. Cleary DW, Jones J, Gladstone RA, Osman KL, Devine VT, Jefferies JM, et al. Changes in serotype prevalence of Streptococcus pneumoniae in Southampton, UK between 2006 and 2018. Sci Rep-Uk. 2022;12(1):13332.

59. Fu J, Yi R, Jiang Y, Xu S, Qin P, Liang Z, et al. Serotype distribution and antimicrobial resistance of Streptococcus pneumoniae causing invasive diseases in China: a meta-analysis. BMC pediatrics. 2019;19:1–9.

60. Choi EH, Kim SH, Eun BW, Kim SJ, Kim NH, Lee J, et al. Streptococcus pneumoniae serotype 19A in children, South Korea. Emerging infectious diseases. 2008;14(2):275.

61. Tin Tin Htar M, Christopoulou D, Schmitt HJ. Pneumococcal serotype evolution in Western Europe. BMC Infect Dis. 2015;15:419.

62. Lo SW, Gladstone RA, van Tonder AJ, Lees JA, du Plessis M, Benisty R, et al. Pneumococcal lineages associated with serotype replacement and antibiotic resistance in childhood invasive pneumococcal disease in the post-PCV13 era: an international whole-genome sequencing study. Lancet Infectious Diseases. 2019;19(7):759–69.

63. Richter SS, Diekema DJ, Heilmann KP, Dohrn CL, Riahi F, Doern GV. Changes in pneumococcal serotypes and antimicrobial resistance after introduction of the 13-valent conjugate vaccine in the United States. Antimicrob Agents Chemother. 2014;58(11):6484–9.

64. Desmet S, Theeten H, Laenen L, Cuypers L, Maes P, Bossuyt W, et al. Characterization of emerging serotype 19A pneumococcal strains in invasive disease and carriage, Belgium. Emerging Infectious Diseases. 2022;28(8):1606.

65. Tiley KS, Ratcliffe H, Voysey M, Jefferies K, Sinclair G, Carr M, et al. Nasopharyngeal Carriage of Pneumococcus in Children in England up to 10 Years After 13-Valent Pneumococcal Conjugate Vaccine Introduction: Persistence of Serotypes 3 and 19A and Emergence of 7C. J Infect Dis. 2023;227(5):610–21.

66. Luck JN, Tettelin H, Orihuela CJ. Sugar-Coated Killer: Serotype 3 Pneumococcal Disease. Front Cell Infect Microbiol. 2020;10:613287.

67. Weinberger DM, Malley R, Lipsitch M. Serotype replacement in disease after pneumococcal vaccination. Lancet. 2011;378(9807):1962-73.

68. Yahiaoui RY, Bootsma HJ, den Heijer CDJ, Pluister GN, Paget WJ, Spreeuwenberg P, et al. Distribution of serotypes and patterns of antimicrobial resistance among commensal in nine European countries. Bmc Infectious Diseases. 2018;18.

69. Brandileone MC, Almeida SCG, Bokermann S, Minamisava R, Berezin EN, Harrison LH, et al. Dynamics of antimicrobial resistance of Streptococcus pneumoniae following PCV10 introduction in Brazil: Nationwide surveillance from 2007 to 2019. Vaccine. 2021;39(23):3207–15.

70. Wu CJ, Lai JF, Huang IW, Shiau YR, Wang HY, Lauderdale TL. Serotype Distribution and Antimicrobial Susceptibility of Streptococcus pneumoniae in Pre- and Post- PCV7/13 Eras, Taiwan, 2002-2018. Front Microbiol. 2020;11:557404.

71. Klugman KP, Black S. Impact of existing vaccines in reducing antibiotic resistance: Primary and secondary effects. Proc Natl Acad Sci U S A. 2018;115(51):12896–901.

72. Løchen A, Croucher NJ, Anderson RM. Divergent serotype replacement trends and increasing diversity in pneumococcal disease in high income settings reduce the benefit of expanding vaccine valency. Sci Rep-Uk. 2020;10(1):18977.

73. Lo SW, Mellor K, Cohen R, Alonso AR, Belman S, Kumar N, et al. Emergence of a multidrug-resistant and virulent Streptococcus pneumoniae lineage mediates serotype replacement after PCV13: an international whole-genome sequencing study. Lancet Microbe. 2022;3(10):e735–e43.

74. Feldgarden M, Brover V, Gonzalez-Escalona N, Frye JG, Haendiges J, Haft DH, et al. AMRFinderPlus and the Reference Gene Catalog facilitate examination of the genomic links among antimicrobial resistance, stress response, and virulence. Sci Rep-Uk. 2021;11(1).

75. Lobb B, Lee MC, McElheny CL, Doi Y, Yahner K, Hoberman A, et al. Genomic classification and antimicrobial resistance profiling of Streptococcus pneumoniae and Haemophilus influenza isolates associated with paediatric otitis media and upper respiratory infection. BMC Infect Dis. 2023;23(1):596.

76. Kapatai G, Sheppard CL, Al-Shahib A, Litt DJ, Underwood AP, Harrison TG, et al. Whole genome sequencing of Streptococcus pneumoniae: development, evaluation and verification of targets for serogroup and serotype prediction using an automated pipeline. PeerJ. 2016;4:e2477.

77. Lee JT, Li X, Hyde C, Liberator PA, Hao L. PfaSTer: a machine learning-powered serotype caller for Streptococcus pneumoniae genomes. Microb Genom. 2023;9(6).

78. Sheppard CL, Manna S, Groves N, Litt DJ, Amin-Chowdhury Z, Bertran M, et al. PneumoKITy: A fast, flexible, specific, and sensitive tool for Streptococcus pneumoniae serotype screening and mixed serotype detection from genome sequence data. Microbial Genomics. 2022;8(12):000904.

79. Hakenbeck R, Bruckner R, Denapaite D, Maurer P. Molecular mechanisms of beta-lactam resistance in Streptococcus pneumoniae. Future Microbiol. 2012;7(3):395–410.

80. Smith AM, Klugman KP. Alterations in MurM, a cell wall muropeptide branching enzyme, increase high-level penicillin and cephalosporin resistance in Streptococcus pneumoniae. Antimicrob Agents Chemother. 2001;45(8):2393–6.

81. Coffey TJ, Daniels M, McDougal LK, Dowson CG, Tenover FC, Spratt BG. Genetic analysis of clinical isolates of Streptococcus pneumoniae with high-level resistance to expanded-spectrum cephalosporins. Antimicrob Agents Chemother. 1995;39(6):1306–13.

82. Grebe T, Hakenbeck R. Penicillin-binding proteins 2b and 2x of Streptococcus pneumoniae are primary resistance determinants for different classes of beta-lactam antibiotics. Antimicrob Agents Chemother. 1996;40(4):829–34.

83. Ribeiro CD, van Roode MY, Haringhuizen GB, Koopmans MP, Claassen E, van de Burgwal LHM. How ownership rights over microorganisms affect infectious disease control and innovation: A root-cause analysis of barriers to data sharing as experienced by key stakeholders. Plos One. 2018;13(5).

84. Kaye J, Heeney C, Hawkins N, de Vries J, Boddington P. Data sharing in genomics--re-shaping scientific practice. Nat Rev Genet. 2009;10(5):331–5.

85. Sabot F. On the importance of metadata when sharing and opening data. BMC Genomic Data. 2022;23(1):79.

86. Schadron T, van den Beld M, Mughini-Gras L, Franz E. Use of whole genome sequencing for surveillance and control of foodborne diseases: status quo and quo vadis. Front Microbiol. 2024;15:1460335.

87. Epping L, van Tonder AJ, Gladstone RA, The Global Pneumococcal Sequencing C, Bentley SD, Page AJ, et al. SeroBA: rapid high-throughput serotyping of Streptococcus pneumoniae from whole genome sequence data. Microb Genom. 2018;4(7).

