## Supplementary material for "Large-scale global molecular epidemiology of antibiotic resistance determinants in *Streptococcus pneumoniae*": Suppl material

Roger Junges, DDS, PhD

Associate Professor

Institute of Oral Biology

Faculty of Dentistry, University of Oslo (UiO)

Postboks 1052 Blindern 0316 - Oslo, Norway

### Supplementary material

*Dataset curation and visualization*

The dataset was cleaned to ensure consistency and readiness for further analysis. All entries were replaced with "Not Available" where data were missing. A new variable "country" was created by extracting country names from the location data. Countries were labeled with their continents using the countrycode package (v1.6.0), distinguishing between North and South America. A "Year" column was added by extracting years from dates. If extraction failed, the original date value was retained. Rows lacking entries in "Complete_Genes" were excluded from further analysis. Gene identifiers were further processed to remove all characters following the "=" delimiter, standardising the gene names. A custom function replaced gene identifiers with their associated resistance classes, generating a new column, "gene_to_class".

We extracted text from the "Location" column, separating the country name from other details by removing everything after the first colon (":"). Terms indicating missing country data (e.g., "not collected", "not available") were identified and replaced with "Not Available" for consistency. The dplyr and stringr packages (v1.5.1) were used to extract the year from the date depending on various formats (e.g., "2006-02-26" to "2006", "2001-09" to "2001"). If neither format was compatible, the initial date value was kept in the "Year" field.

To extract and analyze genes marked as "=COMPLETE" from the AMR.genotypes column, a function called extract_complete_genes was created to find and extract all gene and combine these into a single string. This function was applied to each row in the AMR.genotypes column, and the results were saved in a new column called "Complete_Genes". Rows where the "Complete_Genes" column was empty were removed. Gene names were further refined by removing text up to the next comma after an "=" sign (e.g., "erm(A)=COMPLETE, pmrA=COMPLETE" to "erm(A),pmrA"). A matrix was created to link genes to their corresponding resistance classes (e.g., "ant(9)-Ib" to "AMINOGLYCOSIDE"). A custom function was defined to search through gene names, and link them with their corresponding resistance class, and store the updated information in a new "gene_to_class" column. Another function was created to list unique resistance classes per genome. Additionally, a function was implemented to identify and extract unique AMR classes for each genome, and the results were stored in the "unique_classes" column. Other variables were created based on the presence or absence of specific genes, resistance to specific classes of antibiotics, and the presence of serotypes included in specific vaccines.

*Serotype classification*

Initial prediction was performed with PneumoKITy [78] and PfaSTer [77]. Sequences that lacked quality or coverage were discarded from the analyses. If the predictions matched, the results were then kept; if the predictions were not a match, we attempted to call each isolate with PneumoCAT [76], which utilizes reads and quality scores enabling the tool to make calls between similar serotypes. To obtain the reads, the biosample accession numbers were tracked through the Sequence Read Archive (SRA), and a pipeline was used to retrieve the SRA IDs, download the sequences, and convert them into fastq format for further processing using NCBI SRA Tool (v3.1.1). If no accession number was available or the fastq sequences were not able to be traced, we searched the NCBI database for information from the authors regarding the serotyping of the deposited isolates. For serogroup 6, user submitted data and SeroBA [87] were employed to call genotype 6E into serotypes 6A or 6B.

*Gene co-occurrence and data visualization*

For gene co-occurrence, we filtered the dataset to include only MDR isolates. We split gene names, created a binary matrix representing gene presence/absence using Base R function, and computed a co-occurrence matrix via matrix multiplication. The resulting matrix was filtered to include only gene pairs over the threshold of 100 occurrences. In addition, we reshaped the matrix into a data frame where each row represents a gene pair and its co-occurrence frequency and visualize a chord diagram with circlize (v0.4.16).

Data visualization was performed using R Studio, with the ggplot2 (v 3.5.1) package used as the primary tool for generating the figures presented in this study. For geospatial analyses and world map creation, the rworldmap (v1.3.8) package was employed. Additionally, heatmaps were constructed using the ComplexHeatmap (v2.18.0) package.

**Table S1.** Serotypes included in each pneumococcal vaccine.

| Vaccine | Serotypes |
|---|---|
| PCV7 | 4, 6B, 9V, 14, 18C, 19F, 23F |
| PCV10 | 4, 6B, 9V, 14, 18C, 19F, 23F, 1, 5, 7F |
| PCV13 | 4, 6B, 9V, 14, 18C, 19F, 23F, 1, 5, 7F, 3, 6A, 19A |
| PCV15 | 4, 6B, 9V, 14, 18C, 19F, 23F, 1, 5, 7F, 3, 6A, 19A, 22F, 33F |
| PCV20 | 4, 6B, 9V, 14, 18C, 19F, 23F, 1, 5, 7F, 3, 6A, 19A, 22F, 33F, 15B, 12F, 11A, 10A, 8 |
| PPSV23 | 4, 6B, 9V, 14, 18C, 19F, 23F, 1, 5, 7F, 3, 19A, 22F, 33F, 15B, 12F, 11A, 10A, 8, 20, 17F, 9N, 2 |
| PCV21 | 1, 3, 4, 5, 6A, 6B, 7F, 9V, 11A, 14, 15A, 15C, 16F, 18C, 19A, 19F, 23A, 23B, 23F, 24F, 31, 35B |

**Table S2.** Distribution of isolates collected by country with MDR score data.

| Country | n | % | Mean MDR score | SEM |
|---|---|---|---|---|
| Argentina | 2 | .0 | 2.00 | 1.00 |
| Australia | 1657 | 2.2 | 0.55 | 0.02 |
| Austria | 1 | .0 | 3.00 | |
| Bangladesh | 94 | .1 | 1.21 | 0.10 |
| Belarus | 72 | .1 | 1.82 | 0.17 |
| Belgium | 213 | .3 | 1.03 | 0.11 |
| Botswana | 1 | .0 | 0.00 | |
| Brazil | 508 | .7 | 1.05 | 0.05 |
| Burkina Faso | 1 | .0 | 2.00 | |
| Cambodia | 56 | .1 | 2.14 | 0.11 |
| Cameroon | 8 | .0 | 1.75 | 0.37 |
| Canada | 1854 | 2.5 | 0.56 | 0.02 |
| Central African Republic | 4 | .0 | 2.00 | 0.41 |
| China | 3166 | 4.2 | 2.75 | 0.01 |
| Colombia | 6 | .0 | 1.50 | 0.67 |
| Croatia | 6 | .0 | 1.50 | 0.34 |
| Czechia | 43 | .1 | 0.70 | 0.14 |
| Denmark | 57 | .1 | 0.46 | 0.13 |
| Egypt | 34 | .0 | 1.91 | 0.18 |
| Ethiopia | 43 | .1 | 1.74 | 0.22 |
| Finland | 1 | .0 | 0.00 | |
| France | 40 | .1 | 2.28 | 0.28 |
| Gambia | 1638 | 2.2 | 0.68 | 0.02 |
| Germany | 330 | .4 | 1.10 | 0.05 |
| Ghana | 55 | .1 | 1.78 | 0.11 |
| Greece | 2 | .0 | 2.50 | 0.50 |
| Greenland | 2 | .0 | 0.00 | 0.00 |
| Hong Kong | 55 | .1 | 1.96 | 0.07 |
| Hungary | 24 | .0 | 0.83 | 0.26 |
| Iceland | 120 | .2 | 3.74 | 0.05 |
| India | 964 | 1.3 | 1.60 | 0.04 |
| Indonesia | 92 | .1 | 1.73 | 0.12 |
| Ireland | 152 | .2 | 0.57 | 0.08 |
| Israel | 1148 | 1.5 | 0.91 | 0.03 |
| Italy | 14 | .0 | 1.43 | 0.25 |
| Japan | 220 | .3 | 2.61 | 0.05 |
| Kazakhstan | 1 | .0 | 1.00 | |
| Kenya | 1 | .0 | 1.00 | |

| | | | | |
|---|---|---|---|---|
| Kuwait | 1 | .0 | 3.00 | |
| Lebanon | 9 | .0 | 2.78 | 0.22 |
| Lithuania | 1 | .0 | 0.00 | |
| Malawi | 4238 | 5.6 | 1.04 | 0.02 |
| Malaysia | 47 | .1 | 1.85 | 0.22 |
| Mexico | 14 | .0 | 3.14 | 0.10 |
| Mongolia | 1 | .0 | 1.00 | |
| Morocco | 42 | .1 | 1.26 | 0.15 |
| Mozambique | 168 | .2 | 0.92 | 0.06 |
| Myanmar | 58 | .1 | 1.60 | 0.15 |
| Nepal | 416 | .6 | 0.79 | 0.05 |
| Netherlands | 1775 | 2.4 | 0.23 | 0.01 |
| New Zealand | 718 | 1.0 | 0.60 | 0.03 |
| Niger | 15 | .0 | 1.27 | 0.23 |
| Norway | 24 | .0 | 0.00 | 0.00 |
| Oman | 1 | .0 | 4.00 | |
| Papua New Guinea | 2 | .0 | 1.00 | 0.00 |
| Peru | 1043 | 1.4 | 1.51 | 0.04 |
| Poland | 284 | .4 | 2.09 | 0.09 |
| Portugal | 192 | .3 | 1.08 | 0.11 |
| Qatar | 95 | .1 | 1.40 | 0.11 |
| Russia | 143 | .2 | 1.12 | 0.11 |
| Senegal | 31 | .0 | 0.94 | 0.19 |
| Singapore | 2 | .0 | 1.50 | 1.50 |
| Slovenia | 91 | .1 | 0.98 | 0.13 |
| South Africa | 5037 | 6.7 | 0.92 | 0.01 |
| South Korea | 45 | .1 | 3.00 | 0.10 |
| Spain | 241 | .3 | 1.29 | 0.08 |
| Sweden | 64 | .1 | 0.14 | 0.07 |
| Switzerland | 11 | .0 | 0.36 | 0.24 |
| Taiwan | 94 | .1 | 3.31 | 0.05 |
| Tanzania | 26 | .0 | 2.04 | 0.23 |
| Thailand | 3072 | 4.1 | 1.65 | 0.02 |
| Togo | 20 | .0 | 1.35 | 0.17 |
| Trinidad and Tobago | 84 | .1 | 0.75 | 0.11 |
| Turkey | 15 | .0 | 2.27 | 0.33 |
| United Kingdom | 1292 | 1.7 | 0.29 | 0.02 |
| United States | 21488 | 28.6 | 0.71 | 0.01 |
| Uruguay | 3 | .0 | 3.00 | 0.00 |
| Vietnam | 18 | .0 | 2.89 | 0.21 |

**Table S3.** Antibiotic resistance genes (ARGs) identified in *S. pneumoniae* isolates.

| Antibiotic class | Gene | Count | Relative to ARGs | Relative to genomes |
|---|---|---|---|---|
| Aminoglycoside | *ant(2")-Ia* | 1 | 0.00 | 0.00 |
| | *ant(6)-Ia* | 4 | 0.00 | 0.01 |
| | *ant(9)-Ib* | 4 | 0.00 | 0.01 |
| | *aph(3')-Ia* | 1 | 0.00 | 0.00 |
| | *aph(3')-IIIa* | 315 | 0.26 | 0.42 |
| | *str* | 18 | 0.01 | 0.02 |

| | | | | |
|---|---|---|---|---|
| | *aac(6')-Ie/aph(2'')-Ia* | 14 | 0.01 | 0.02 |
| | *aadD1* | 2 | 0.00 | 0.00 |
| Beta-lactam | *bla2* | 1 | 0.00 | 0.00 |
| | *blaARL* | 3 | 0.00 | 0.00 |
| | *blaARL-2* | 11 | 0.01 | 0.01 |
| | *blaI* | 20 | 0.02 | 0.03 |
| | *blaPC1* | 11 | 0.01 | 0.01 |
| | *blaR1* | 23 | 0.02 | 0.03 |
| | *blaTEM* | 3 | 0.00 | 0.00 |
| | *blaTEM-1* | 1 | 0.00 | 0.00 |
| | *blaTEM-116* | 4 | 0.00 | 0.01 |
| | *blaTEM-135* | 1 | 0.00 | 0.00 |
| | *blaTEM-171* | 1 | 0.00 | 0.00 |
| | *blaZ* | 13 | 0.01 | 0.02 |
| | *mecA* | 17 | 0.01 | 0.02 |
| | *mecA1* | 5 | 0.00 | 0.01 |
| | *mecI* | 2 | 0.00 | 0.00 |
| | *mecR1* | 2 | 0.00 | 0.00 |
| | divergent-*pbp1a*s | 14436 | 11.77 | 19.21 |
| | divergent-*pbp2b*s | 25690 | 20.94 | 34.18 |
| | divergent-*pbp2x*s | 24183 | 19.71 | 32.17 |
| Bleomycin | *bleO* | 1 | 0.00 | 0.00 |
| | *ble-Sh* | 1 | 0.00 | 0.00 |
| Fosfomycin | *fosB* | 5 | 0.00 | 0.01 |
| | *fosY* | 1 | 0.00 | 0.00 |
| Fusidane | *fusD* | 10 | 0.01 | 0.01 |
| Glycopeptide | *vanC1* | 1 | 0.00 | 0.00 |
| | *vanG* | 1 | 0.00 | 0.00 |
| | *vanR-C* | 1 | 0.00 | 0.00 |
| | *vanS-C* | 1 | 0.00 | 0.00 |
| | *vanT-C* | 1 | 0.00 | 0.00 |
| | *vanXY* | 1 | 0.00 | 0.00 |
| | *vanXY-C* | 1 | 0.00 | 0.00 |
| Lincosamide | *lnu(A)* | 35 | 0.03 | 0.05 |
| | *lnu(A)'* | 3 | 0.00 | 0.00 |
| | *lsa(C)* | 10 | 0.01 | 0.01 |
| | *sal(A)* | 1 | 0.00 | 0.00 |
| | *vga(A)-LC* | 5 | 0.00 | 0.01 |
| Macrolide | *erm(A)* | 4 | 0.00 | 0.01 |
| | *erm(B)* | 10525 | 8.58 | 14.00 |
| | *erm(C)* | 41 | 0.03 | 0.05 |
| | *erm(X)* | 3 | 0.00 | 0.00 |
| | *mef(A)* | 12032 | 9.81 | 16.01 |
| | *mph(C)* | 11 | 0.01 | 0.01 |

| | | | | |
|---|---|---|---|---|
| | *msr(A)* | 5 | 0.00 | 0.01 |
| | *msr(D)* | 12020 | 9.80 | 15.99 |
| Phenicol | *catA* | 2833 | 2.31 | 3.77 |
| | *catA1* | 3 | 0.00 | 0.00 |
| | *catA16* | 145 | 0.12 | 0.19 |
| | *catP* | 1 | 0.00 | 0.00 |
| | *cmx* | 4 | 0.00 | 0.01 |
| Streptothricin | *sat4* | 209 | 0.17 | 0.28 |
| Sulfonamide | *sul1* | 3 | 0.00 | 0.00 |
| Tetracycline | *tet(32)* | 160 | 0.13 | 0.21 |
| | *tet(38)* | 2 | 0.00 | 0.00 |
| | *tet(K)* | 132 | 0.11 | 0.18 |
| | *tet(L)* | 16 | 0.01 | 0.02 |
| | *tet(M)* | 19429 | 15.84 | 25.85 |
| | *tet(O)* | 33 | 0.03 | 0.04 |
| | *tet(S)* | 150 | 0.12 | 0.20 |
| | *tet(W)* | 2 | 0.00 | 0.00 |
| | *tetA(60)* | 1 | 0.00 | 0.00 |
| | *tetB(60)* | 1 | 0.00 | 0.00 |
| Trimethoprim | *dfrE* | 3 | 0.00 | 0.00 |
| | *dfrG* | 28 | 0.02 | 0.04 |
| | *dfrS1* | 12 | 0.01 | 0.02 |
| **Total** | | **122673** | | |

**Table S4.** Distribution of macrolide resistance in *S. pneumoniae* determined by *erm(B)* and/or *mef(A)/msr(D)* across continents between 2015-2022.

| | *mef(A)* | | *erm(B)* | | Dual genotype | | Total genomes |
|---|---|---|---|---|---|---|---|
| | n | % | n | % | n | % | |
| Africa | 482 | 16.07 % | 37 | 1.23 % | 13 | 0.43 % | 3000 |
| Asia | 652 | 35.45 % | 1451 | 78.90 % | 456 | 24.80 % | 1839 |
| Europe | 70 | 6.36 % | 158 | 14.36 % | 49 | 4.46 % | 1102 |
| North America | 3997 | 21.49 % | 1826 | 9.82 % | 331 | 1.78 % | 18595 |
| South America | 155 | 40.90 % | 161 | 42.48 % | 77 | 20.32 % | 379 |
| Oceania | 52 | 3.40 % | 136 | 8.89 % | 19 | 1.24 % | 1529 |
| Total | 5409 | 20.42 % | 3786 | 14.29 % | 945 | 3.58 % | 26495 |

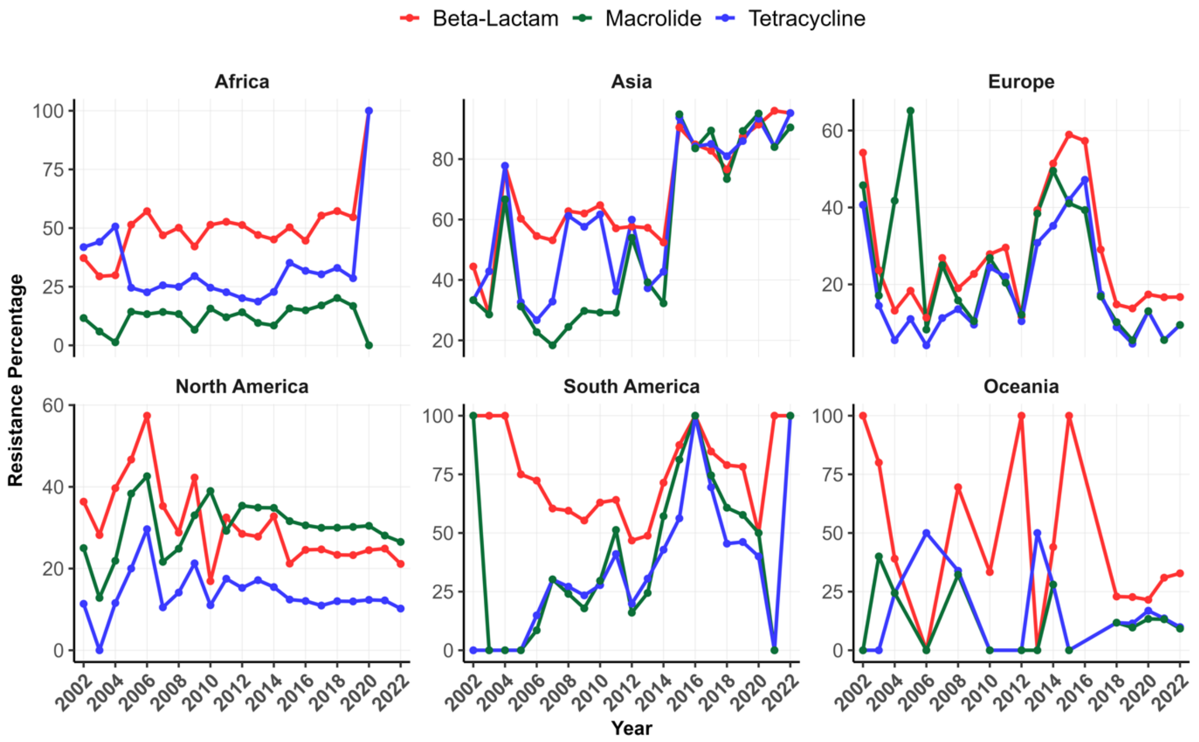


**Fig S1**. Distribution of resistance to the three most common antibiotic classes (beta-lactams, macrolides, and tetracyclines) over time. Each segment of the panel represents a continent.

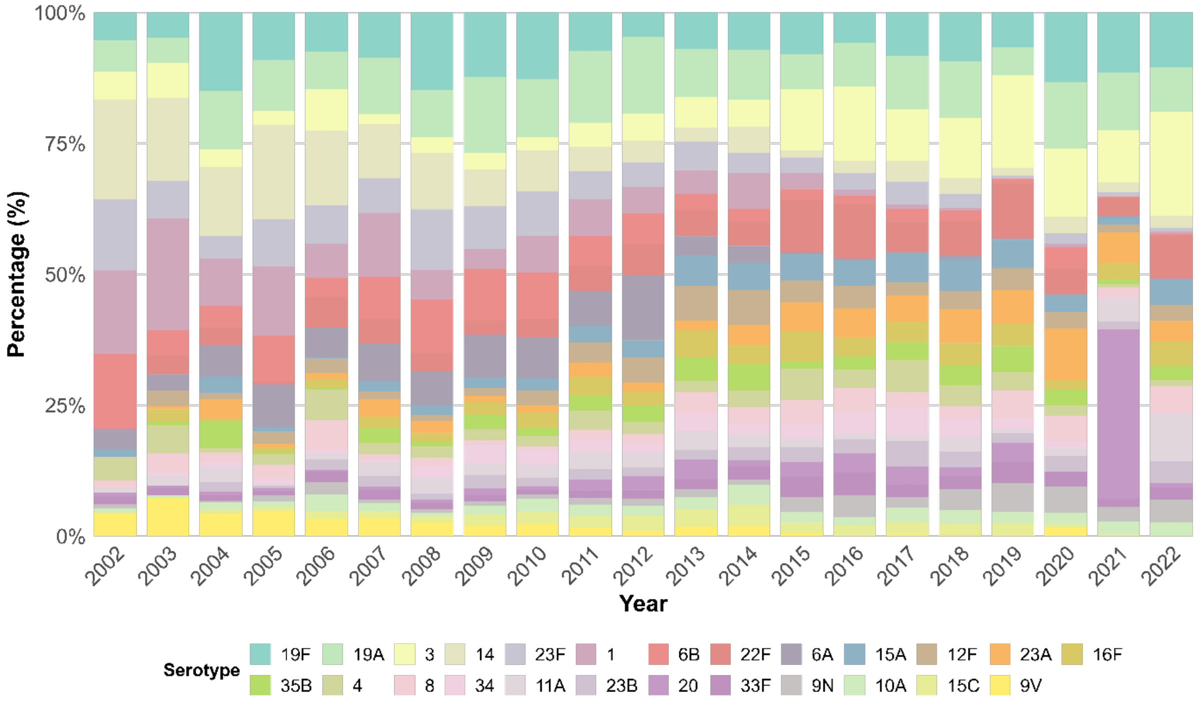


**Fig S2.** Stacked bar chart showing the serotype composition each year. The 25 most prevalent serotypes are included in the figure.

**Table S5.** Distribution and MDR rate of serotypes with more than 100 genomes available and their trends of development. Serotypes whose trend was not statistically significant were considered stable. **p<0.01***p<.001

| Serotype | Number of genomes | % of MDR | Year estimate | Trend | Statistically significant |
|---|---|---|---|---|---|
| **19F** | 3792 | 63.2 | -0.03 | Down | *** |
| **19A** | 3224 | 34.8 | 0,03 | Up | *** |
| **3** | 2604 | 8.1 | -0,01 | Stable | --- |
| **14** | 2160 | 24.4 | 0.04 | Up | *** |
| **23F** | 2036 | 43.3 | 0.03 | Up | *** |
| **1** | 1837 | 0.8 | 0.05 | Up | ** |
| **6B** | 1821 | 42.5 | 0.03 | Up | *** |
| **22F** | 1724 | 1.6 | 0.06 | Up | *** |
| **6A** | 1533 | 5.6 | 0.02 | Stable | --- |
| **15A** | 1043 | 43.3 | 0.03 | Up | *** |
| **12F** | 1028 | 8.6 | -0.01 | Stable | --- |
| **23A** | 1012 | 28.4 | 0.08 | Up | *** |
| **16F** | 948 | 2.5 | -0.02 | Stable | --- |
| **35B** | 909 | 1.8 | 0.06 | Up | *** |
| **4** | 893 | 1.6 | 0.03 | Stable | --- |
| **8** | 770 | 0.4 | 0.01 | Stable | --- |
| **34** | 701 | 3.1 | -0.06 | Down | ** |
| **11A** | 670 | 0.9 | 0.06 | Up | *** |
| **20** | 638 | 2 | -0.04 | Stable | --- |
| **23B** | 638 | 0.8 | 0.04 | Up | ** |
| **33F** | 637 | 1.9 | 0.07 | Up | *** |
| **9N** | 613 | 0.3 | 0.02 | Stable | --- |
| **10A** | 595 | 4.5 | 0.06 | Up | *** |
| **15C** | 598 | 12.9 | 0.06 | Up | *** |
| **9V** | 559 | 7.9 | 0.07 | Up | *** |
| **5** | 485 | 1.9 | 0.01 | Stable | --- |
| **13** | 480 | 11.7 | 0.13 | Up | *** |
| **21** | 447 | 2.7 | -0.15 | Down | *** |
| **35F** | 445 | 0.4 | 0.01 | Stable | --- |
| **17F** | 434 | 3.9 | 0.08 | Up | *** |
| **15B** | 426 | 6.9 | 0.00 | Stable | --- |
| **6C** | 412 | 7.5 | 0.02 | Stable | --- |
| **7C** | 382 | 6.3 | -0.03 | Stable | --- |
| **7A** | 378 | 0.5 | 0.03 | Stable | --- |
| **31** | 335 | 0.9 | 0.05 | Stable | --- |
| **18C** | 300 | 0.67 | 0.07 | Stable | --- |
| **38** | 295 | 2.4 | -0.14 | Down | ** |
| **7F** | 242 | 0 | 0.09 | Stable | --- |
| **6D** | 194 | 3.7 | 0.11 | Up | *** |
| **10B** | 167 | 2.9 | 0.13 | Up | *** |
| **18A** | 112 | 3.6 | 0.02 | Stable | --- |

19F

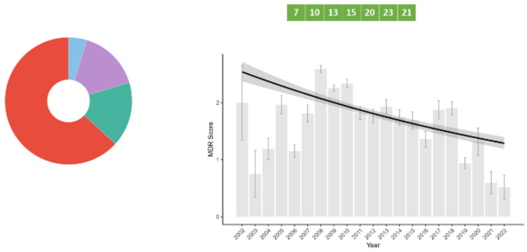


19A

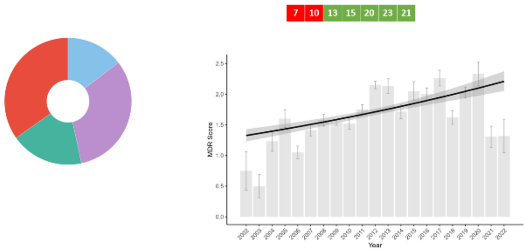


3

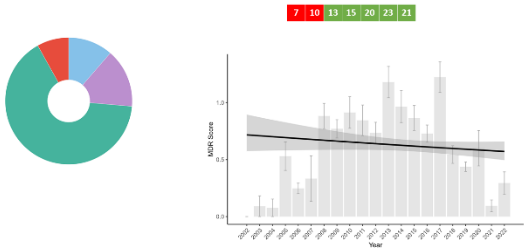


14

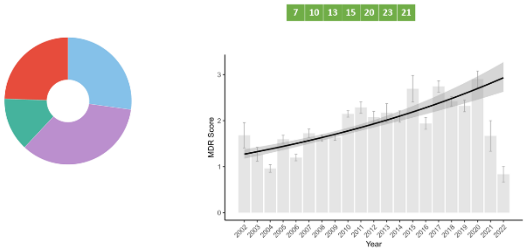


23F

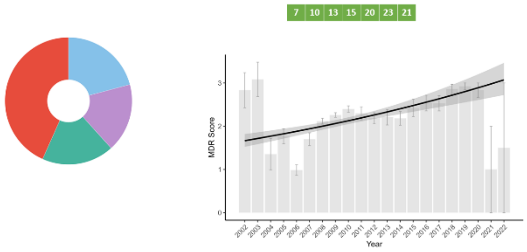


10A

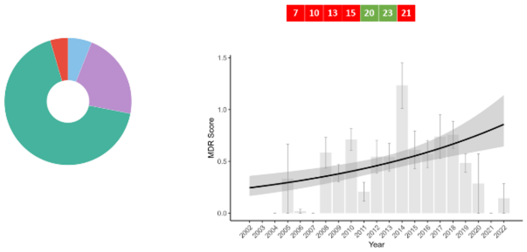


6B

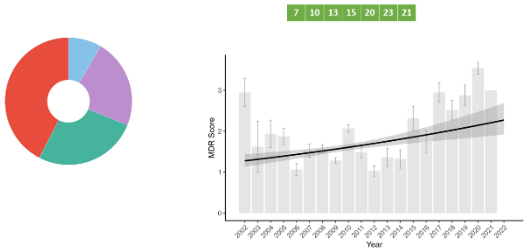


22F

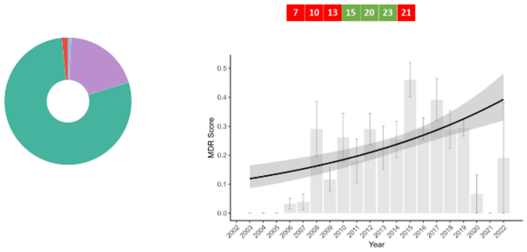


6A

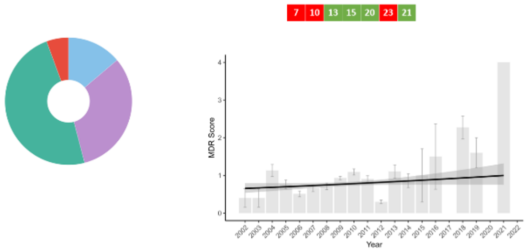


15A

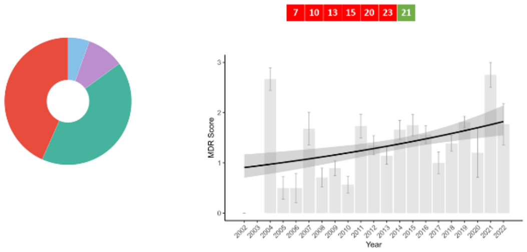


12F

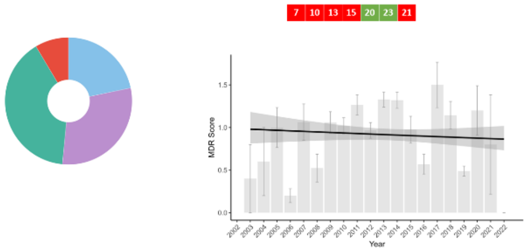


23A

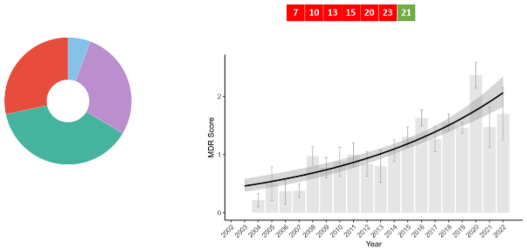

16F

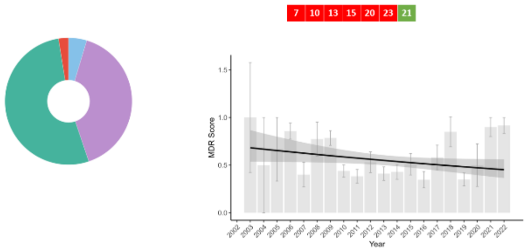


35B

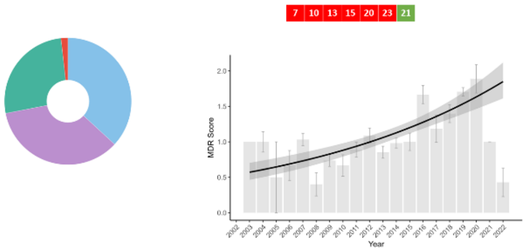


4

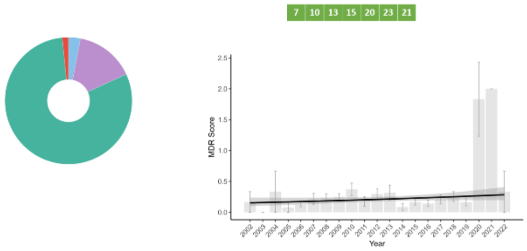


8

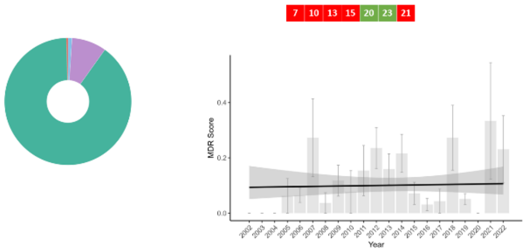


34

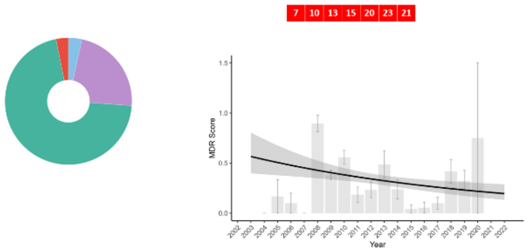


11A

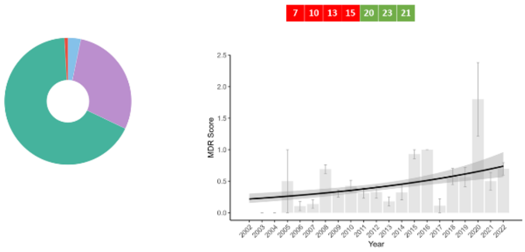


20

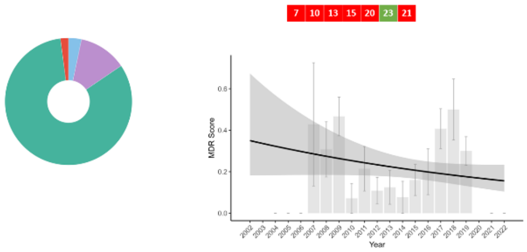


23B

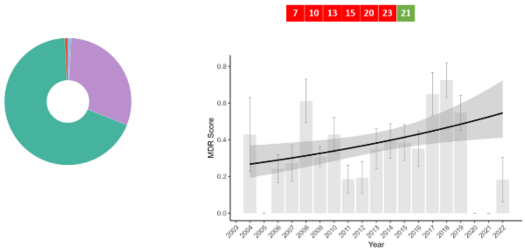


33F

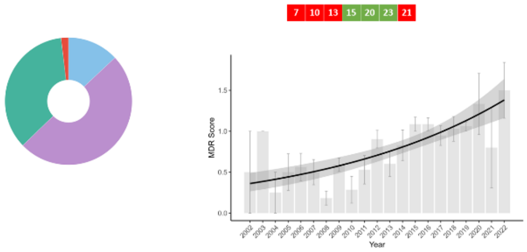


9N

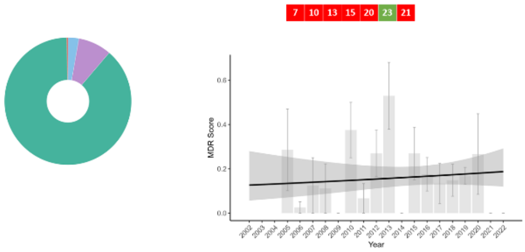


10A

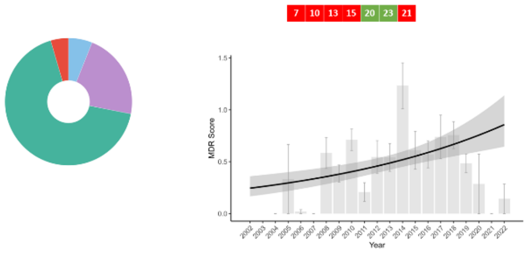


15C

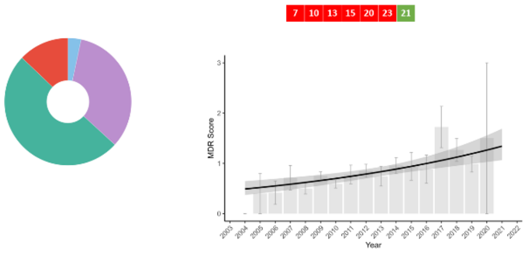

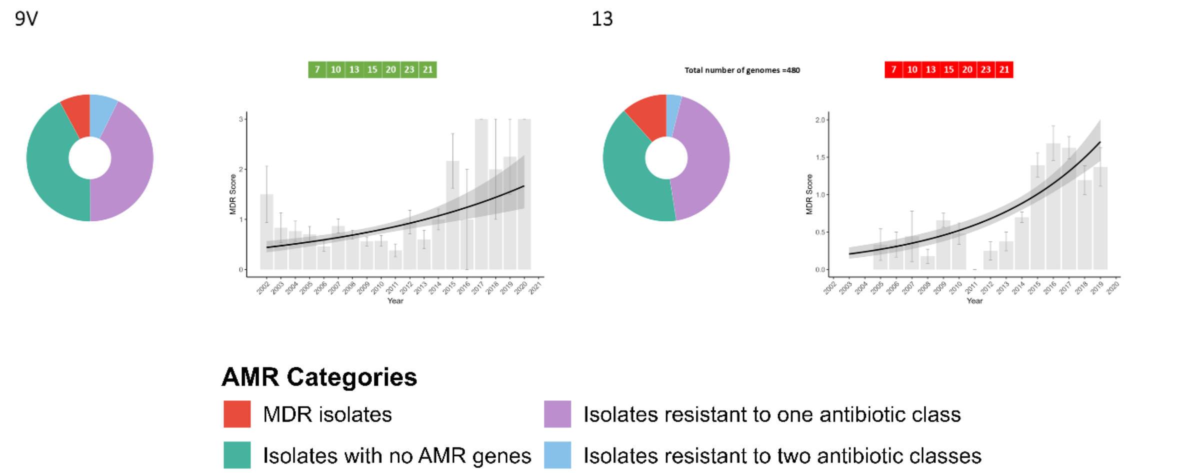


**Fig S3.** Trends over time for individual serotypes and fitted lines for negative binomial regression. Red sections in the donut plot indicate MDR, while green indicates resistance to 2 antibiotic classes, and purple indicates resistance to 1 antibiotic class. Graphs to the right show average MDR score with SEM for every year from 2002 to 2022. Green blocks in the top right indicate inclusion in the vaccine whereas red blocks indicate non-inclusion.

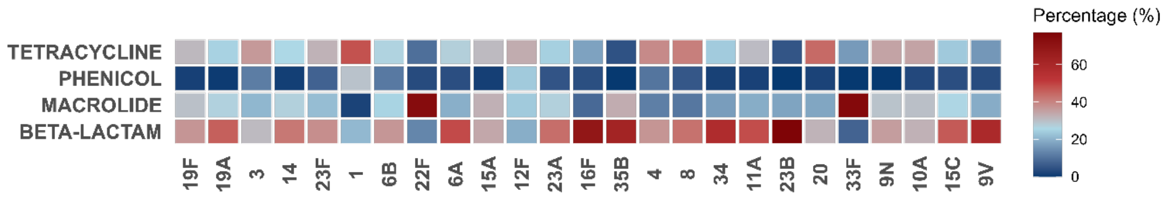


**Fig S4.** Reported resistance to the four most common antibiotic classes in the 25 serotypes with most genomes. Proportions are shown as the percentage of serotype resistance to the corresponding antibiotic class.

**Table S6.** MDR rate of serotypes with over 100 genomes that have not been included in any vaccine formulations.

| Serotype | MDR | not MDR | Total | MDR% |
| --- | --- | --- | --- | --- |
| **34** | 22 | 679 | 701 | 3.1 |
| **13** | 56 | 424 | 480 | 11.7 |
| **21** | 12 | 435 | 447 | 2.7 |
| **35F** | 2 | 443 | 445 | 0.4 |
| **6C** | 31 | 382 | 413 | 7.5 |
| **7C** | 24 | 358 | 382 | 6.3 |
| **7A** | 2 | 376 | 378 | 0.5 |
| **38** | 7 | 288 | 295 | 2.4 |
| **6D** | 7 | 187 | 194 | 3.6 |
| **10B** | 5 | 162 | 167 | 3 |
| **18A** | 4 | 108 | 112 | 3.6 |
| **22A** | 3 | 102 | 105 | 2.9 |
